# Determinants of cysteine coverage by chemoproteomics

**DOI:** 10.64898/2026.10.12.758458

**Authors:** Ioannis G. Riziotis, Harry Wilders, Andrea Cairoli, Hugues Lambert, Jamie Dark, George Biggs, Emma E. Cawood, William J. McCarthy, Luke Nightingale, Paula Magarinos, Mike Gavrielides, Karen Ambrose, Amy Strange, Mark Skehel, Simon J. Boulton, James Lumley, David House, Katrin Rittinger, Jacob Bush

## Abstract

Covalent chemoproteomics has become a powerful route to expanding the druggable proteome, enabling proteome-wide screening of compounds that covalently modify nucleophilic residues, most commonly cysteine. Yet typical experiments detect only 5–18% of the ∼250,000 cysteines encoded by the human proteome, and the determinants of this limited coverage remain poorly defined. Here we combine cysteine featurisation, explainable machine learning (ML) and robust statistical filtering to establish what governs cysteine detectability in iodoacetamide-desthiobiotin (IA-DTB) chemoproteomics. Using a scalable feature-generation pipeline, we computed physicochemical and structural properties for cysteines and the corresponding tryptic peptides and trained models that predict detectability with high accuracy. Shapley analysis identifies protein abundance and intrinsic peptide properties governing LC–MS behaviour as the dominant determinants, whereas cysteine structural descriptors contribute comparatively little, indicating that detection is constrained downstream of probe reactivity. We further introduce a distance-based statistical filter that improves reproducibility across replicates, and show that cell line selection, subcellular fractionation and alternative electrophiles together expand coverage by >30%. More broadly, this work moves optimisation of chemoproteomics workflows from empirical trial-and-error towards rational, data-driven design.

**Graphical abstract:** 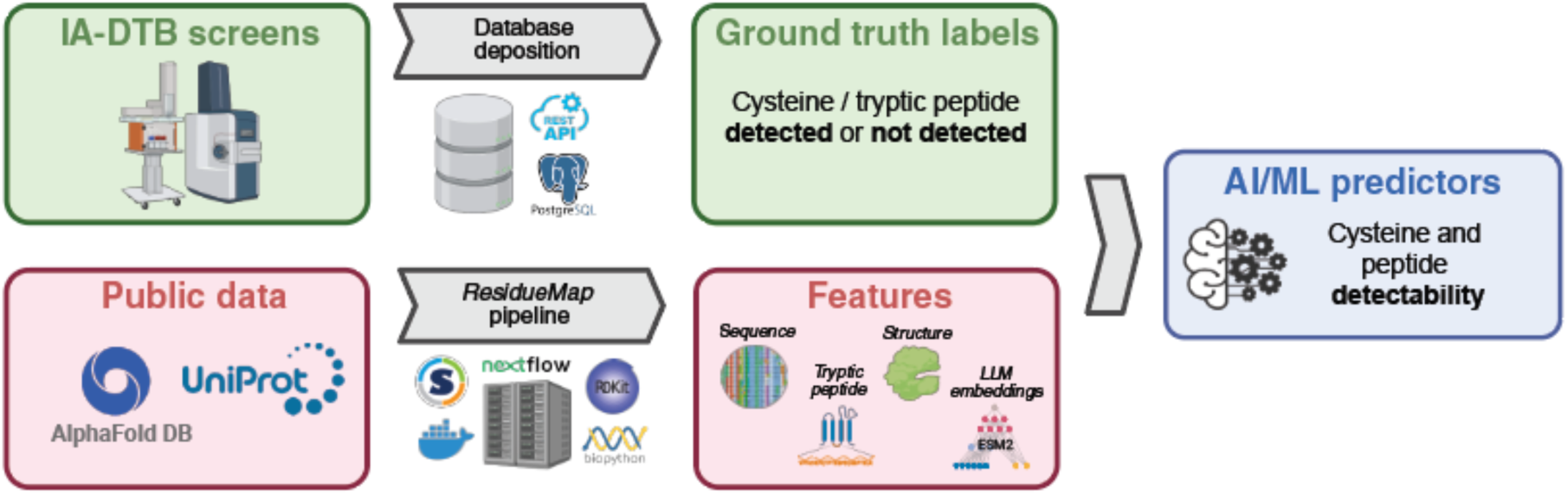

## Introduction

Insights from ‘omics technologies are transforming our understanding of biology and disease, yet for ∼90% of proteins we have no drug-like modulators to validate these insights and potentially progress towards the clinic^1^. Covalency has emerged as a powerful approach to target proteins that have been assumed to be undruggable. Covalent ligands bond irreversibly with nucleophilic amino acids, most prominently cysteines, which can deliver high potency and enable binding to shallow or cryptic pockets, disordered regions, and interfaces involved in protein-protein interactions^2^. Several covalent inhibitors have reached clinical approval, including neratinib, a rationally designed inhibitor of the Tyr-kinase domain of HER1 for breast cancer^3^ and more recently sotorasib, which targets KRAS^4^.

Mass spectrometry-based chemoproteomic screening of covalent ligands, pioneered by Cravatt et al.^5^, has become an essential tool for covalent drug discovery, providing a proteome-wide approach to identifying reactive and ligandable cysteine residues in native environments^6^. By combining electrophilic cysteine-targeting probes that are conjugated with a pull-down tag, (e.g. iodoacetamide-desthobiotin (IA-DTB) for cysteine targeting and streptavidin binding), with proteomic Mass Spectrometry (LC-MS), cysteine reactivity can be interrogated at proteome scale. This approach is particularly effective for protein regions inaccessible to purified protein-based screening, including disordered regions and pockets at protein-protein interfaces, as well as insoluble and high-molecular weight proteins.

Despite the widespread adoption, covalent chemoproteomics still faces two key challenges: i) comprehensive coverage of accessible cysteine sites across the proteome, and ii) robust detection of peptides, meaning peptides are reproducibly detected with consistent intensity^7^. There are ∼250,000 cysteine residues encoded by the human proteome, of which typical chemoproteomics experiments detect 15,000-40,000; this represents just 5-18% of the cysteinome^8^. Understanding which cysteines are detected, as well as the determinants of detectability, is therefore essential to both deploy these technologies for expansion of the druggable proteome and to assess target engagement of emerging covalent leads and candidates.

Addressing these challenges requires predictive approaches to understand peptide behaviour and improve covalent chemoproteomics workflows. Previous work has explored the relationship between cysteine exposure and detectability using iodoacetamide probes, finding poor correlation with pPSE^9^. Other studies have explored peptide detectability in global proteomics experiments, without the cysteine enrichment step^10–14^. These have illustrated the importance of peptide hydrophobicity, charge, and length in determining MS ionisation and fragmentation efficiency^15^. Additionally, predictive models for MS detectability have emphasised the role of amino acid composition in driving ionisation behaviour^14^. However, no studies have pulled together all possible determinants of detectability for cysteine enrichment chemoproteomics, which is essential to understanding the scope and limitations of the technology.

With this study we aim to advance chemoproteomics research by integrating detailed peptide characterisation pipelines, explainable ML, and robust statistical methodologies to address the challenges of detectability in MS workflows. By elucidating the molecular factors influencing it, our approach serves as a foundation for improving chemoproteomics platforms towards better proteome coverage, expanding our understanding of peptide behaviour in MS experiments and ultimately accelerating target discovery,

## Results

### Chemoproteomics dataset

To assess the determinants of cysteine coverage by chemoproteomics, we analysed a dataset in which IA-DTB had been used to enrich peptides from HEK293T cell lysates (as previously reported^8^). This is the representative dataset used throughout all downstream analyses of this study. Our experimental protocol involved cysteine labelling using iodoacetamide desthiobiotin (IA-DTB), followed by trypsin digestion, streptavidin enrichment and analysis of captured peptides using a timsTOF Pro2 mass spectrometer coupled to an Evosep One liquid chromatography system^16^ (Figure 1). To establish a robust hybrid spectral library with broad peptide coverage, we pooled lysates from three diverse cell types: HEK293T, Jurkat, and eHAP (see Biggs, Cawood et al.^8^, and Methods). Merging these distinct proteomic backgrounds creates a composite reference that enhances identification across varied samples.

**Figure 1:**
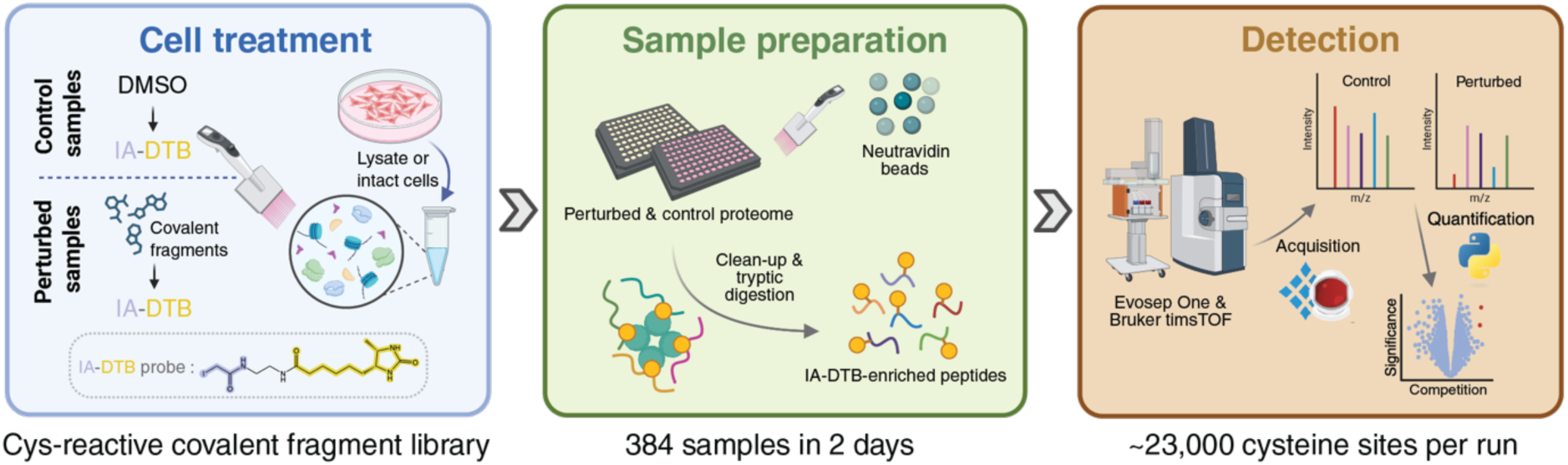
Experimental framework used in this work. Overview of the label-free chemoproteomics platform that was used to generate ground truth data for proteins that have had cysteine residues modified by covalent fragments, using a highly reactive IA-DTB probe. Created with BioRender.com

### Statistical identification of robustly detected peptides

A key step in cysteine profiling by chemoproteomics is filtering of the data for those peptides that are considered robustly detected: defined here as being quantified with sufficient reproducibility and confidence to be included in downstream statistical analyses without introducing high levels of noise. Conventionally, filters are employed by defining empirical thresholds to intensity, data completeness (DC) and CV, which, however, can introduce overly rigid boundaries that exclude peptides that strongly meet two criteria but narrowly miss a third. To balance maximal cysteinome coverage while maintaining high data reproducibility, we developed a statistical filter to identify robustly detected peptides. Our approach compares each peptide to an “ideal peptide” characterised by maximal intensity and DC, and minimal CV within the same experiment. We normalise these three metrics and compute the Euclidean distance to this ideal (Figure 2a). An optimal distance cutoff is then determined by fitting two-component Gaussian mixture to the distance distribution; taking the intersection of these components as the robustness threshold, ensuring maximum inclusion of robustly detected peptides, while minimising noise (Figure 2b).

**Figure 2:**
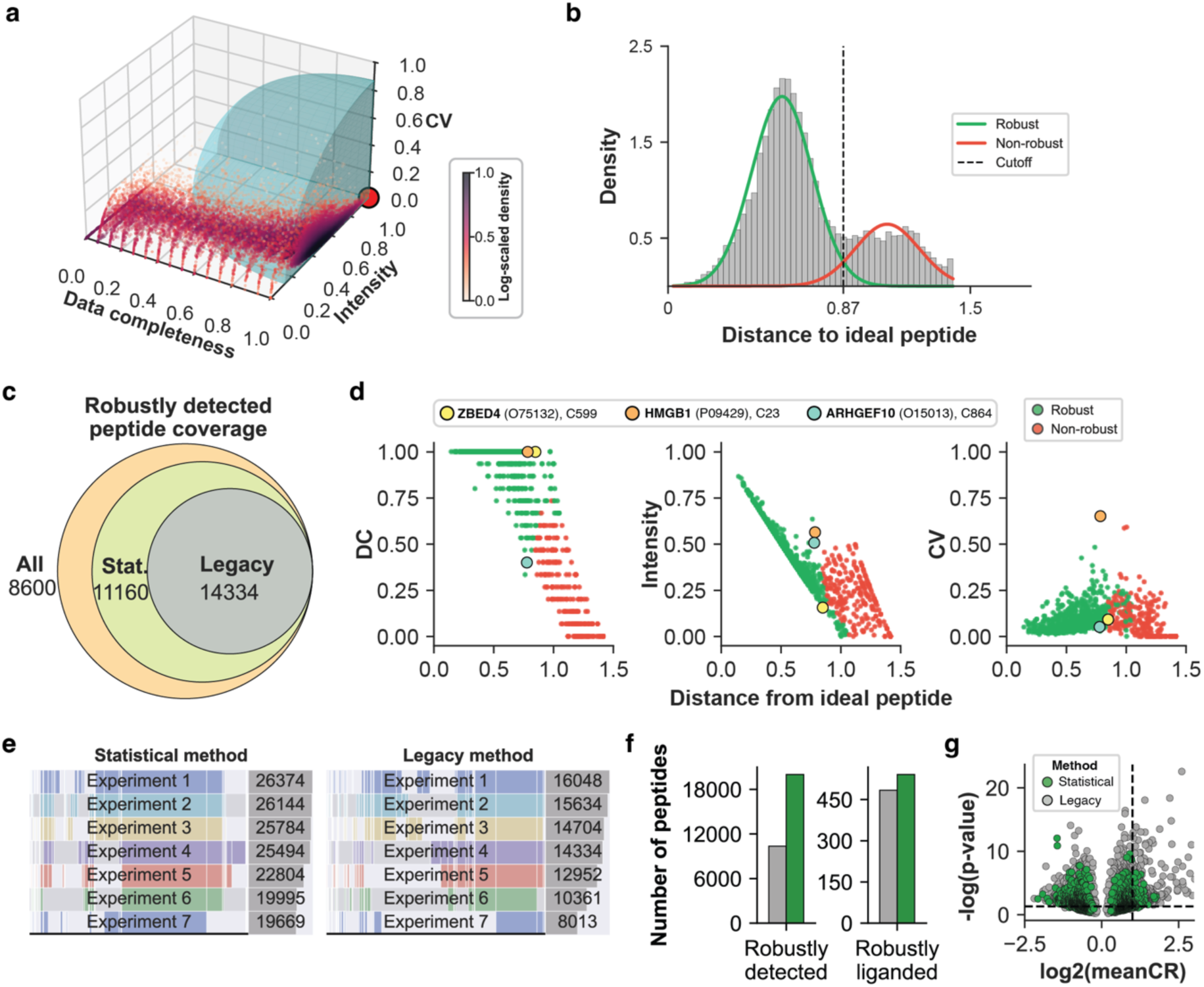
Robust detection of cysteine sites with MS-based label-free chemoproteomics. a) 3D landscape of detected DMSO-treated peptides in terms of quality metrics: data completeness (DC), mean intensity and coefficient of variation of intensity (CV) across replicates. Each point represents a detected peptide with metrics scaled to 0-1 range. A red circle indicates the “ideal peptide” of maximum DC and Intensity, and minimum CV. A blue spherical area on the 3D maps designates the zone of robustly detected peptides, with its radius (distance cutoff) calculated as the intersection points of the two gaussians fitted on the distance distributions. b) Distribution of distances of each peptide to the ideal with a bimodal mixture of gaussians shown as coloured curves. c) Coverage of robustly detected peptides over all detected peptides, when applying the legacy metric-threshold based filtering method, and the statistical filtering method described here. d) Quality metrics of a random sample of 1000 peptides over distance to the ideal peptide. Representative examples of peptides defined as robustly detected by the algorithm are highlighted as distinctly coloured dots. e) Validation of reproducibility of statistical robustness filtering vs legacy filtering, over 7 independent IA-DTB experiments. Experiment coverage and intersection of robustly detected peptides is shown as a supervenn plot, where each horizontal bar represents the set of robustly detected peptides in a single experiment, and its segments indicate the subsets of robustly detected peptides shared with other experiments. The total number of robustly detected peptides is shown in the gray boxes on the right. f) Comparison of statistical and legacy robustness filtering in terms of reflection of robustly detected peptides to robustly liganded peptides, in a single IA-DTB experiment of covalent fragment screening. g) Volcano plot of robustly liganded cysteine sites from the same IA-DTB experiment as panel f, when applying the statistical and legacy robustness filtering methods. The x-axis shows the affinity of a liganding event expressed as the log2-transformed competition ratio (CR) and the y-axis is the significance, expressed as the -log10-transformed p-value across four replicates. p-values were calculated with Welch’s unpaired t-test. Dashed lines indicate an empirical significance threshold to identify interactions as hits, specifically p-value < 0.05 and a CR > 2.

Overall, this method delivered 25,494 robustly detected peptides (23,154 unique cysteines), a significant increase in the proportion compared to using defined metric thresholds (intensity ≥ 100, DC ≥ 0.75, CV ≤ 0.4, defined as legacy filtering) (Figure 2c). Example cysteine sites illustrate the method’s advantages (Figure 2d): ZBED4_C599 (low Intensity) and HMGB1_C23 (high CV) would be excluded by defined thresholds yet are retained by the statistical filter; conversely, AP5Z1_C301 has high intensity and low CV despite DC below the hard cutoff and is also retained. For validation, we applied the method on seven independent IA-DTB experiments on HEK293T cell lysates. The statistical filter produced more consistent cysteinome coverage and reduced batch effects compared to using defined thresholds (Figure 2e). Additionally, we tested how the increased number of robustly detected peptides impacted outcomes in a competitive profiling experiment. Eight fragments carrying a chloroacetamide electrophile group were screened in HEK293T cell lysates, and conventional and statistical filtering were applied to DMSO controls. The statistical method nearly doubled the number of robustly detected peptides, and also gave a 12% increase in the number of liganded cysteines, from 484 to 541 ((≥2-fold change, -log(p) ≥1.3, Figures 2e,f). The proportional change in identified hits provided confidence that the inclusion of additional peptides had not introduced a significant increase in noise / false positives.

### Analysis of protein and peptide coverage

Having compiled a dataset of robustly detected peptides from HEK293T cells, we analysed proteome coverage from the perspective of protein and peptide properties. In total we identified 32,823 cysteine-containing peptides from 7,888 proteins (Figure 3a). In the human proteome, there are predicted to be 20,577 proteins and 262,515 cysteines, thus, while good coverage of proteins was obtained (∼40%), coverage of cysteines was much lower (∼12%). The key features expected to influence cysteine detectability were i) protein abundance, and ii) occurrence of cysteines in tryptic peptides of length compatible with MS protocols. The influence of protein abundance was investigated using intensity values from global proteomics experiments, as reported in ProteomicsDB^16^. Many proteins (∼28% of proteome) are reported as not detectable in global proteomics, and these were typically also not detected in cysteinome profiling, confirming that abundance is still a key determinant of cysteine detectability. For those proteins that were detected in global proteomics, only ∼60% were also detected in our cysteinome profiling, and protein intensity in global proteomics showed little influence on detectability in cysteinome profiling (Figure 3b). Other features that were explored include analysis of the number of cysteines per protein, revealing that more cysteine residues render a protein more detectable (Figure 3c). Also, the coverage of proteins according to cell localisation was analysed by enrichment analysis of Gene Ontology (GO) terms for detected / undetected proteins, revealing, not unexpectedly, that transmembrane and secreted terms were strongly enriched among undetected proteins (Figure 3d). However, GO cell localisation terms are overlapping, so do not provide a suitable criterion for more deterministic analyses.

**Figure 3:**
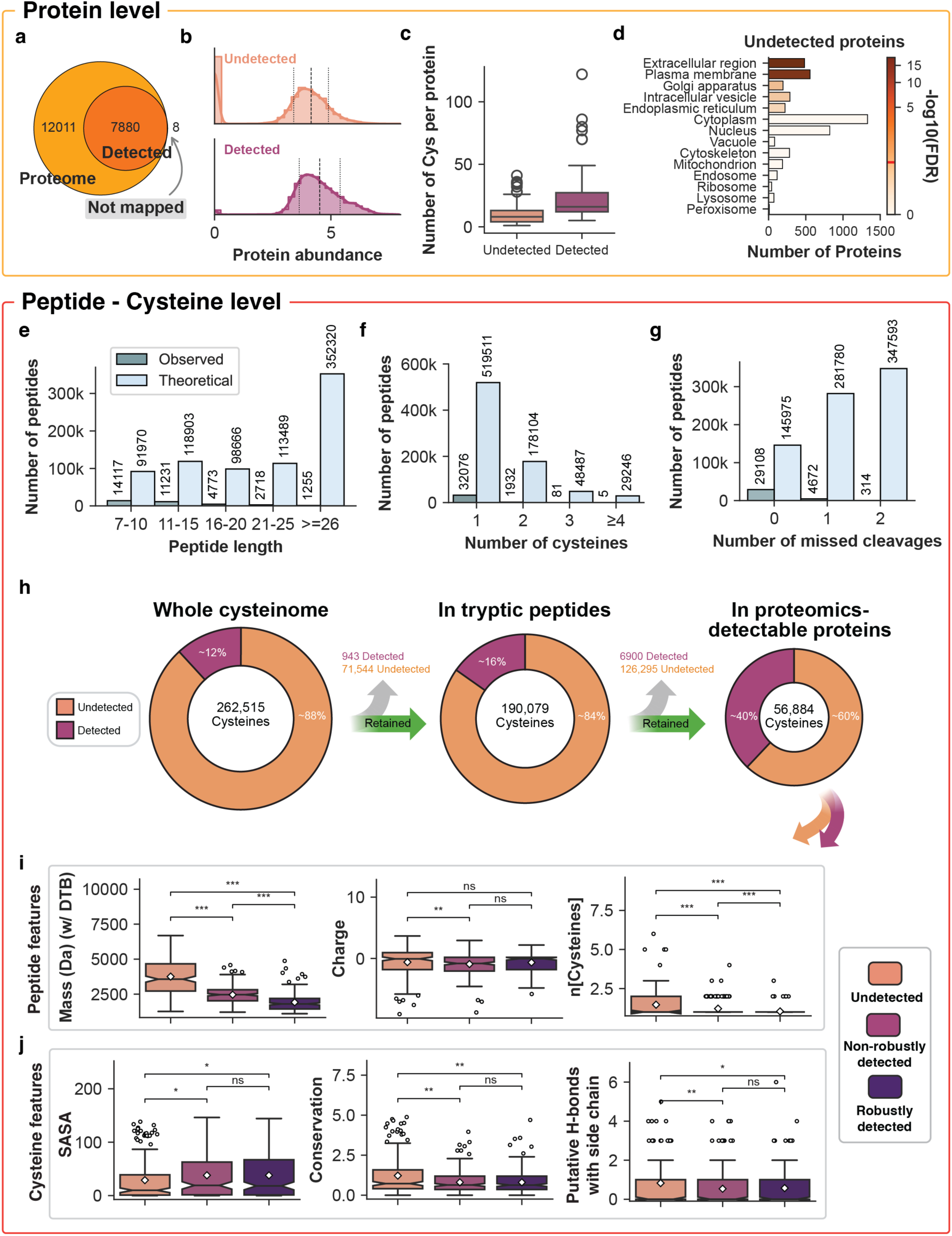
Analysis of ground truth chemoproteomics data. a) Proteome coverage in IA-DTB screening of HEK293T cell lysate, in terms of number of proteins. b) Distributions of average protein abundance across known cell lines (from publicly available global proteomics data) in undetected and detected proteins from the IA-DTB experiment. Vertical lines indicate the mean (centre) and standard deviation (outer lines) of the distributions. c) Distribution of cysteine counts per protein stratified by protein detectability. Proteins are classified as "Detected" if at least 5 cysteines are detected, otherwise "Undetected". Boxplots show median, quartiles, and outliers for 100 randomly sampled proteins from each category d) GO terms-based cellular localisation enrichment analysis of proteins undetected by the IA-DTB workflow in HEK-293T cells, compared to the maximum detectable proteome in the spectral library. Enrichment significance is coloured by p-value, and the absolute number of proteins in each cellular compartment is shown on the x-axis. A red mark on the colour bar indicates the 0.05 FDR significance threshold. e-g) Distributions of basic properties of detected DTB-enriched peptides in HEK293T cell lysates (observed), and the in silico digested proteome (theoretical). h) Filtering process of ground truth dataset for downstream feature trends analysis. i-j) Boxplots comparing peptide (i) and cysteine (j) feature values in undetected, non-robustly detected and robustly detected peptides. Asterisks denote statistically significant differences between detectability groups calculated by Mann-Whitney statistical testing (p < 0.05: *, p < 0.01: **, p < 0.001: **, non-significant: ns). White diamonds indicate feature mean values. For cysteine structural features analysis, data was further filtered to exclude non-pocketed cysteines and cysteines engaged in disulfide bonds.

We next explored proteome coverage at the peptide level. Mass spectrometry protocols are typically optimised for tryptic peptides of length 7-52 residues, which gives a theoretical coverage of 83.7% of cysteines (missed cleavages, N ≤ 2) or 72.8% of cysteines (missed cleavages, N=0) (Figure 3e-g). Our experimental dataset contained predominantly perfectly cleaved peptides (∼82%), therefore, analyses were only conducted on those (N=0), to avoid introducing many theoretical but rarely observed miscleaved peptides (N=1 and N=2). Combining the two key filters, abundance and presence in tryptic peptide, we estimated that the number of cysteines expected to be detectable, would be 56,884, of which 30,369 were detected in HEK293T (23,154 robustly detected), which is a large improvement compared to 12% coverage of the full theoretical proteome (Figure 3h).

### Peptide and residue features

A scalable computational pipeline – *ResidueMap* – was developed to calculate a broad set of parameters that might influence cysteine detectability. *ResidueMap* outputs the following types of features for any given set of UniProt sequences^17^: 1) physicochemical descriptors for tryptic peptides derived from adjustable *in silico* tryptic digestion. These include, amongst others, mass, length, charge, TPSA, MolMR and hydrophobicity; 2) structural properties of cysteine (or lysine, serine and tyrosine) residues and their environment in the context of the protein in tertiary structure, calculated on AlphaFold models^18,19^ using established and bespoke algorithms for estimating p*K*_a_^20^, solvent-accessible surface area (SASA) ^21^, interatomic interactions and other parameters; 3) Protein abundance values as averaged MS2 intensities from multiple proteomics experiments deposited in ProteomicsDB^16^. A detailed breakdown of calculated features can be found in Methods; the pipeline was implemented with Nextflow^22^ and is available as open-source software at https://github.com/FrancisCrickInstitute/ResidueMap. We also calculated deep learning embeddings of whole protein sequences, tryptic peptides, and cysteines, using the ESM-2 Protein Language Model^23^. These were generated by *ESMembedder*, a Python pipeline also available as open source software at https://github.com/FrancisCrickInstitute/esm_embedder.

### Features affecting peptide and cysteine detection

To further investigate the features that influence detectability among these 57,199 theoretically detectable peptides, we analysed differences across two distinct feature categories using Mann-Whitney U testing (*p* < 0.05:∗, *p* < 0.01:∗∗, *p* < 0.001:∗∗∗) (Figures 3i–j). Specifically, we evaluated global peptide-level traits (mass, charge and number of cysteines) separately from local cysteine microenvironment features (solvent accessibility [SASA], conservation and number of putative H-bonds involving Cys side chains). This distinction reflects the underlying biochemistry: the local 3D structural environment determines cysteine reactivity with IA-DTB, directly governing labelling efficiency and, downstream, whether the parent peptide is enriched and detected. These trends indicate that peptide mass exerts the strongest influence on detectability, whereas local structural exposure of the cysteine residue plays a minimal role. However, while this statistical approach revealed clear patterns, inferring direct causality remains non-trivial due to strong feature correlations (Figures S2–S5).

### Predicting Cys-containing peptide detectability

While the numerical trends of key features provided some insights into the determinants of whether proteins and peptides are detected, none provided definitive criteria of detection, and it was anticipated that detectability is determined by a complex combination of factors. ML-based analyses provide a deeper understanding of feature importance and interactions, addressing these complexities and offering more insight. To explore this, ML models were built to predict which theoretical Cys-containing peptides are detected based on the cysteine annotations using protein, peptide and cysteine features.

The cysteinome data was split into test and train sets as described in Methods. We initially employed 5 models: linear regression, logistic regression, random forest, SVM, and CatBoost (Table 1). Models were built to predict the probability of whether the cysteine-containing peptide can be robustly detected in HEK293T; they were trained on a balanced training set and evaluated on the protein held-out split. The balanced test view contained 8,672 data points with approximately equal class representation. The natural-prevalence view restores all filtered points for the same 2,691 held-out proteins and contains 35,894 points, of which 4,293 are positive (12.0% prevalence). On the balanced test view, CatBoost was the strongest model in this comparison (AUROC = 0.9652) (Figure 4a). On the natural-prevalence test view, CatBoost also performs best (AUROC = 0.9633, AUPR = 0.7780, F1 = 0.6504). AUROC remains similar across the two class distributions because it measures ranking, while AUPR and F1 respond to prevalence and the chosen 0.5 threshold. Optimisation of this threshold is expected to lead to higher F1 scores.

**Figure 4:**
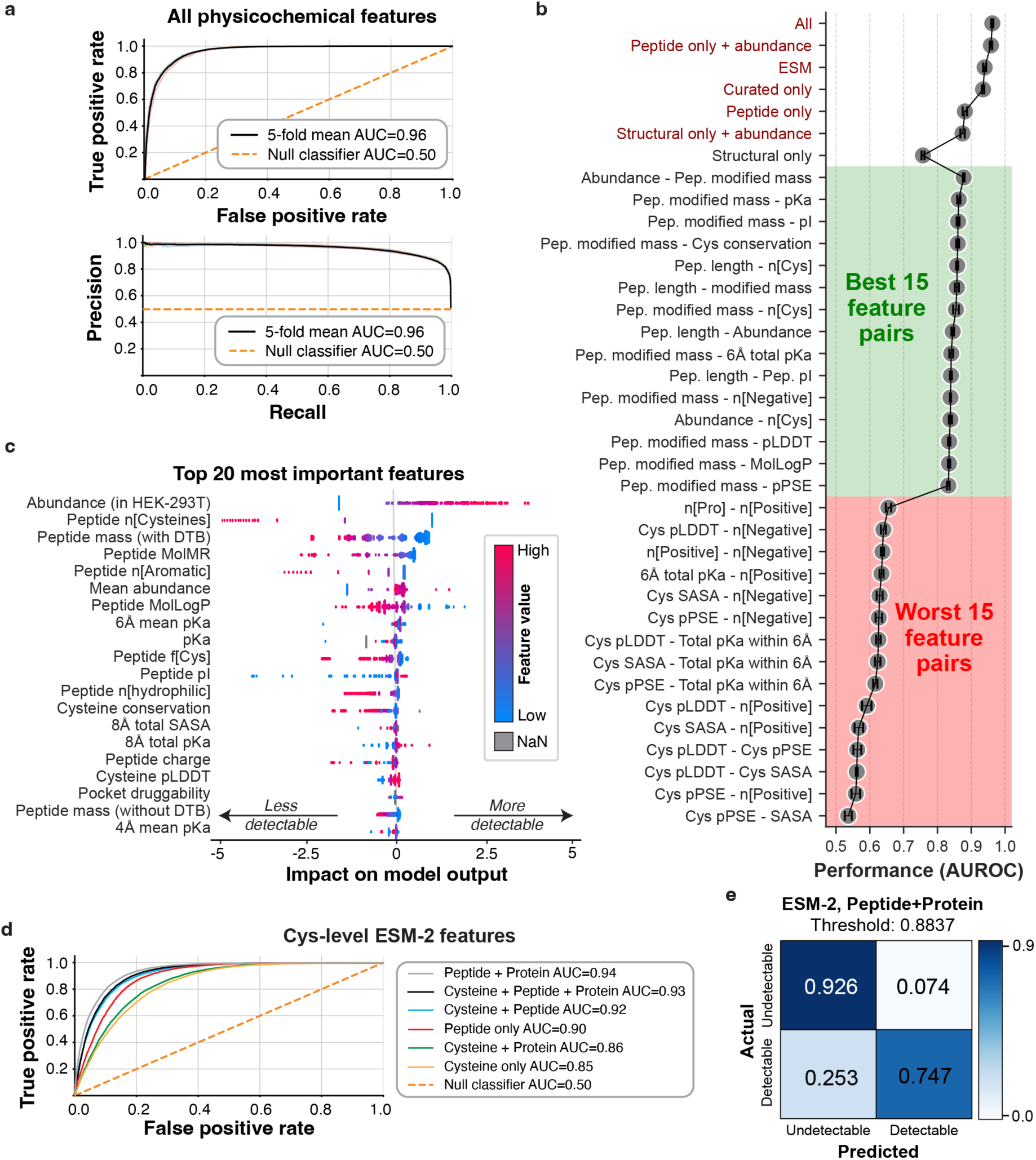
Machine learning to predict and understand Cys-containing peptide detectability in MS-based chemoproteomics. a) ROC and precision-recall (PR) curves (top and bottom panel respectively) of best-performing CatBoost model for predicting robust peptide MS detectability. b) Comparison of CatBoost model performance, expressed as the area under the receiver operating characteristic curve (AUROC), when different combinations of features are used. For detailed lists of features, refer to Methods. c) Ranking of the top 20 features predicting peptide detectability as calculated by explainable ML (GlassBox model). Feature importance is shown as a “beeswarm” plot, where the x-axis shows the Shapley value (impact on model decision) of each feature in the y-axis. Colour of each point corresponds to the value of the feature, effectively showing how trends of features influence the model decision (more positive Shapley value --> more detectable, more negative Shapley value --> more undetectable. d) Detectability ROC curves for different ESM-2 embedding combinations, when training and validating an mlp model on the human cysteinome. e) Row-normalised confusion matrix of best-performing ESM-2-based classifier at optimized decision threshold.

**Table 1:** Machine learning model implementations for detectability prediction.

| Model | Balanced test set |  |  | Unbalanced test set |  |  |
| --- | --- | --- | --- | --- | --- | --- |
|  | AUROC | AUPR | F1 | AUROC | AUPR | F1 |
| Linear Regression | 0.9508 | 0.9452 | 0.8744 | 0.9257 | 0.6158 | 0.4147 |
| Logistic Regression | 0.9605 | 0.9535 | 0.8938 | 0.9312 | 0.6587 | 0.4885 |
| Random Forest | 0.9603 | 0.9486 | 0.8937 | 0.9201 | 0.5405 | 0.5090 |
| SVM | 0.9608 | 0.9534 | 0.8999 | 0.9315 | 0.6346 | 0.5011 |
| <u>CatBoost</u> | <u>0.9652</u> | <u>0.9589</u> | <u>0.9018</u> | <u>0.9633</u> | <u>0.7780</u> | <u>0.6504</u> |

To determine the most important features in obtaining model accuracy, CatBoost was used with different combinations features (Figure 4b). Using only peptide features and protein abundance led to equally high AUROC values, while in contrast, using structural features, even with the addition of abundance data, perform poorly; indicating it is the peptide physicochemical properties (such as ionizability) that are more important in governing detection than IA-DTB reactivity. We subsequently explored how well models performed when using just two features, revealing again that peptide features scored highly, while structural features lead to poor performance.

We subsequently applied an explainable ML framework (GlassBox) and ranked the top 20 features contributing to peptide detectability (Figure 4c). Ranking was calculated via Shapley analysis, which quantifies the contribution of each feature to a model’s prediction by distributing the impact of each feature value in every prediction. The top-ranking features included protein abundance, peptide mass, peptide length, and physicochemical properties such as hydrophobicity, charge, aromaticity, and polarity. The most important features are the protein abundance in the cell line and the peptide mass. In addition, several features correlated to mass such as length and TPSA appear important, as well as features related to the ease of ionisation such as the peptide pI (isoelectric point) and MolMR (a metric of polarisability). These findings are consistent with previous reports from PeptideRanger^24^ where pI, molecular weight and features correlated with it were highlighted as important. Again, structural features do not rank high, however, we do see p*K*_a_ of cysteine and its environment in the top 10 features, likely reflecting that cysteine reactivity enhances the binding of IA-DTB, although this appears to be less significant than the peptide ionisability.

We also explored how peptide and protein embeddings based only on sequence could be used to predict detectability, using ESM-2 embeddings. ESM embeddings are high-dimensional descriptor vectors of each residue in a protein derived by using a protein LLM (esm2_t36_3B_UR50D, a general-purpose model) trained on a large set of protein sequences^25^. We tested different architectures of cysteinome embedding, including protein, cysteine and peptide level embeddings in various combinations and merging strategies (see methods for details and Figure S6), using a multi-layer perceptron neural network (mlp) architecture^26^. We found that these embeddings provide excellent prediction performance, with a maximum AUROC of 0.94 when concatenating the whole sequence embedding vector with the vector of the shortest tryptic peptide carrying the cysteine of interest (Figure 4d). This is interesting as it suggests that in the absence of peptide or abundance knowledge, the whole-protein and tryptic peptide deep learning embedding can effectively encapsulate those properties critical for detection. Finally, a user-friendly cysteine detectability predictor was constructed using the optimised top-performing model (concatenated peptide and protein embeddings), in which the decision threshold was calibrated to maximise balanced accuracy (Figure 4e) on an unbalanced test set. The open-source code and application are available under a GNU-GPL 3.0 license at https://github.com/FrancisCrickInstitute/cysteine-detectability-predictor.

### Experimental approaches to expand the detectable cysteinome

Drawing upon the above insights into the determinants of cysteine detectability, we designed a series of experiments aimed at improving the coverage of the human cysteinome beyond the 23,154 robustly detected cysteines detected in the HEK293T cell system. Three strategies were explored: i) differential protein expression, ii) cellular localisation, and iii) diverse electrophilic reactivities and physicochemical properties of the DTB probe.

Our initial strategy focused on identifying optimal cell lines to maximise incremental cysteine coverage. We posited that variations in global protein expression across different cell types would enable the detection of cysteines from proteins of lower abundance in HEK293T cells (Figure 5a)^27^. Principal Component Analysis (PCA) was applied to global proteomics-derived protein expression profiles^16,28^ from a panel of commonly used human cell lines, focusing on the expression diversity of 13,646 proteins undetected in HEK293T cells (Figure 5b). This analysis revealed that while most detectable proteins remain constant across cell lines (Figure S7), there is significant protein abundance variability, particularly between cancer (e.g., Jurkat) and embryonic (e.g., HES-3) cell lines. To validate this finding experimentally, we performed IA-DTB screening of seven widely available cell lines, chosen for their high expression diversity of proteins not detected in HEK293T. This indeed yielded gains in coverage, with Jurkat cells yielding the most significant expansion (Figure 5c). While the incremental improvement per cell line was small (∼5%), across all 7 cell lines we achieved 28% increase in coverage, highlighting that judicious selection of cells lines for screening can help ensure good coverage of targets of interest. To further evaluate whether combining cell lines could increase coverage, we analysed IA-DTB-treated mixed lysates^29^ from pairwise combinations of these eight cell lines. Pairing cell lines produced a moderate but significant increase in proteome coverage compared with individual cell lines (median 34.5% versus 32.8%; *p* = 7.27 × 10^−4^), with the magnitude of the gain varying across cell-line pairs (Figure S10a,b).

**Figure 5:**
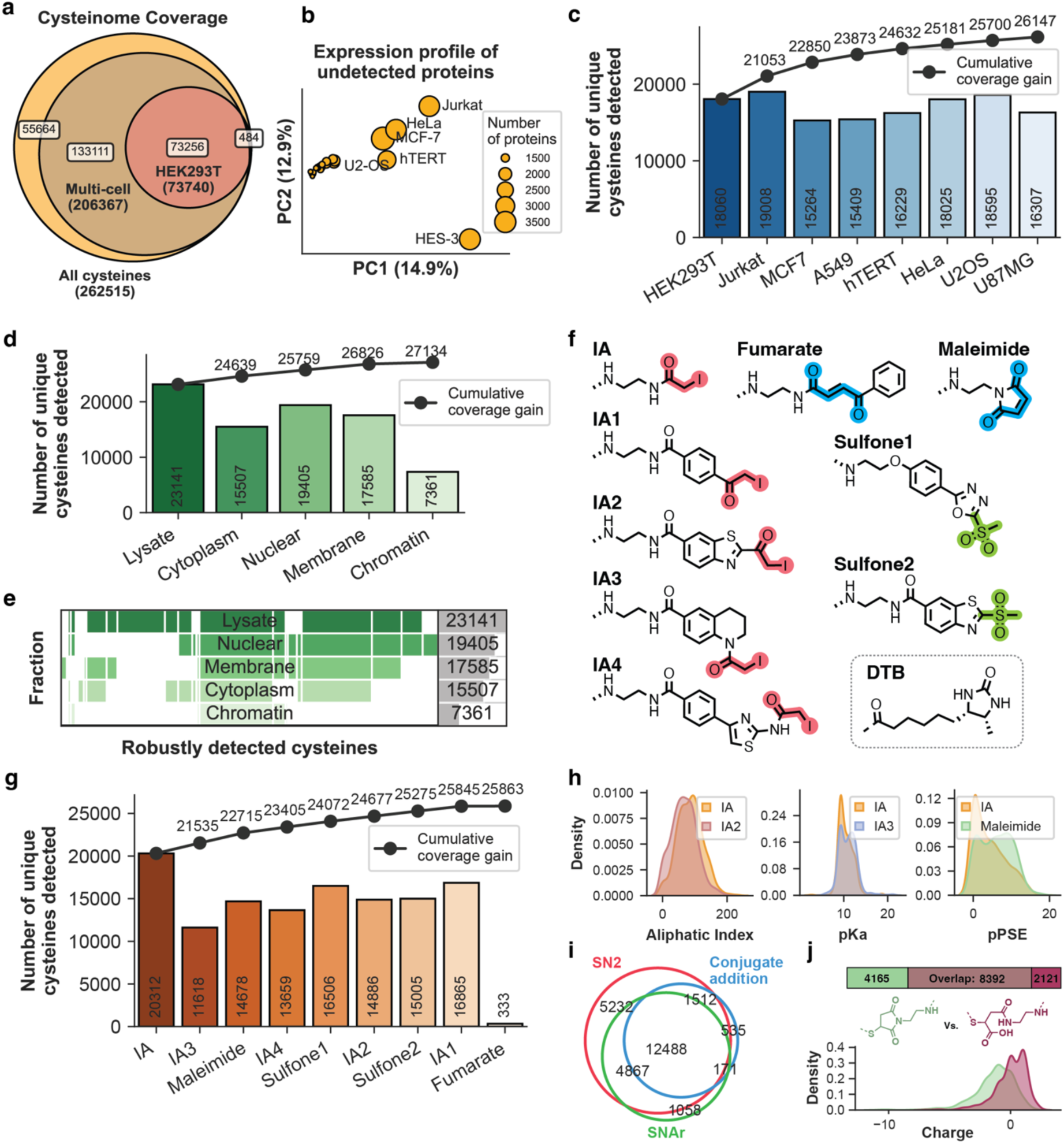
Experimental methods for the expansion of the detectable cysteinome. a) Venn diagram showing the human cysteinome that is detectable by global proteomics, in our HEK293T reference cell line and when combining expression data from multiple cell lines obtained from public resources. b) PCA map showing the difference in expression profile of 13646 proteins that could not be detected by the IA-DTB workflow in HEK293T cells. Each circle is a human cell line, sized based on the number of expressed proteins that were not detected in HEK293T cells. Larger distance between circle indicates higher difference in protein expression profile. Percentage of variance for each PC is shown in parentheses. c) Cysteinome coverage by IA-DTB in alternative cell lines. d) Cysteinome coverage by IA-DTB when a cellular fractionation assay is applied to the cell lysate. e) Cysteinome coverage overlap among cellular fractions. Total cysteine counts are shown on the right. f) Library of alternative electrophile probes. The moiety for covalent attachment is highlighted in different colours. g) Cysteinome coverage by IA-DTB when alternative probes are used. h) Representative feature distributions in samples of 500 peptides (aliphatic index) or cysteines (pKa and pPSE) uniquely detected by pairs of electrophilic probes. i) Cysteinome coverage overlap among probes attaching covalently with different chemistries (S_n_2: IA, IA1, IA2, IA3; Conjugate Addition: Fumarate, Maleimide; S_n_Ar: Sulfone1, Sulfone2). j) Cysteinome coverage (top) and charge distribution of 500 peptides uniquely detected by unhydrolysed (green) and hydrolysed (magenta) forms of Maleimide probe.

Recognising that membrane and secreted proteins are underrepresented in standard whole-cell lysates (Figure 3b), we also investigated subcellular fractionation to increase coverage of cysteinome in specific compartments. We tested the effects of subcellular fractionation of HEK293T lysate in nuclear, cytosolic, membrane and chromatin fractions (Figure 5d,e). This approach yielded a total increase of 3,314 cysteines compared to the unfractionated lysate (+11%). Most of these newly detected cysteines were associated with membrane-bound and nuclear proteins that were previously less detectable under standard lysis conditions.

Finally, we developed and tested a panel of nine electrophile probes designed to modulate the mass, charge, and lipophilicity of labelled peptides, with the aim of capturing peptides with a broader range of physicochemical properties and varied cysteine reactivities. This panel included probes with *α,β*-unsaturated carbonyls and heteroaromatic sulfones, representing different mechanisms of electrophilicity (Figure 5f). Across the tested probes, a significant collective increase in cysteinome coverage was observed, identifying an additional 5,311 cysteines (+25% coverage). While the original IA-DTB probe consistently afforded the greatest individual coverage (Figure 5g), the alternative probes provided complementary detection, with different chemistries showing preference for different peptide chemical properties and cysteine structural environments (Figures 5h, S8 and S9). Probe IA2, which introduced additional hydrophobicity via a phenyl ring, detected peptides with a compensatory higher degree of hydrophilicity. Probes IA3 and Maleimide, incorporating distinct structural scaffolding, were hypothesised to impose different steric interactions within protein pockets, leading to increasing coverage of more reactive (IA3) and buried (Maleimide) cysteines respectively (Figure 5h). The Fumarate probe yielded only 367 enriched peptides, likely due to steric hindrance preventing streptavidin enrichment. We next investigated whether the covalent bonding mechanism of the electrophile affected cysteinome coverage. The nine tested probes were grouped into three distinct mechanisms: S_N_2 (IA-DTB, IA1-4), conjugate addition (Fumarate and Maleimide) and S_N_Ar (Sulfone1-2). Of the 26,352 unique cysteines robustly detected across the probes, 19,113 (72.5%) showed no mechanism selectivity (detected by ≥2 probe mechanisms). Among the mechanistically selective cysteines, the majority (5,780 cysteines) were selective towards S_N_2 mechanisms (represented by IA-DTB, IA1-4), contrasting with fewer selective detections for conjugate addition (305 cysteines) and S_N_Ar (1,154 cysteines) (Figure 5i). Interestingly, partial ring-opening hydrolysis of Maleimide was observed, with both the intact and hydrolysed products detected on modified peptides. The hydrolysed product, containing an acidic functionality, was found to impact peptide detection, enriching for more positively charged peptides compared to the non-hydrolysed adduct (Figure 5j).

## Discussion

Here, we have developed an integrated methodology that combines peptide characterisation, explainable machine learning, and a robust statistical filtering approach to predict and interpret the detectability of cysteine-containing peptides in IA-DTB DIA chemoproteomics^8^ (Figure 1). This platform enables efficient profiling of cysteines for covalent compound screening, but its achievable cysteinome coverage is shaped by both biology (e.g. expression) and analytical constraints (e.g. MS detectability). By coupling ground-truth IA-DTB detection labels with a broad feature set spanning protein abundance, peptide physicochemical properties, and cysteine microenvironment descriptors, we aimed to provide both predictive performance and mechanistic insight that can guide rational workflow optimisation. While ML methods have been applied to proteomics in the past, for example, in predicting peptide retention time^15^ and ionisability^11^, to our knowledge this is the first study to integrate explainable ML methodologies with cysteine-specific peptide detectability analysis.

To facilitate our analyses, we utilised extensive feature calculation for peptides and cysteines. This comprehensive feature library aims to capture the nuanced peptide and cysteine properties that might affect MS detectability. A central finding of this work is that IA-DTB “detectability” is driven primarily by protein abundance and intrinsic peptide properties that govern LC–MS behaviour (e.g., peptide mass/length and features related to ionisation/fragmentation), whereas cysteine structural descriptors contribute comparatively little. Across both univariate comparisons and ML models, peptide-level features, especially mass and related correlates, dominate the separability between detected and undetected peptides, while structural exposure metrics show only weak effects. Consistent with this, models trained using peptide features plus abundance achieved near-maximal performance, whereas models using structural features (even when combined with abundance) performed substantially worse, and structural-only models performed poorly. Although cysteine microenvironment properties such as p*K*_a_ appear among informative features, their impact is secondary relative to abundance and peptide ionisability-related properties. This is crucial for interpretation of IA-DTB datasets: IA-DTB profiling is often implicitly assumed to preferentially report the most reactive and/or most surface-exposed cysteines, yet our analyses indicate that enrichment/reactivity effects are frequently not the primary bottleneck. On the contrary, detection is largely constrained downstream by whether the resulting labelled tryptic peptide is MS-favourable and derived from a sufficiently abundant protein.

It was also striking to observe how strongly protein language model embeddings performed for detectability prediction. Using ESM-2 sequence embeddings, we achieved high predictive accuracy (with best performance obtained by combining whole-protein and tryptic-peptide embeddings), despite not explicitly providing the model with experimentally measured abundance or handcrafted physicochemical descriptors such as lipophilicity. This supports the idea that deep sequence representations can act as rich, “compressed” descriptors that implicitly capture multiple determinants of detectability (e.g., compositional patterns related to peptide ionisation, digestibility proxies, and other sequence-encoded correlates), offering an attractive alternative when structural models or reliable abundance estimates are unavailable.

Beyond identifying determinants, our study also helps clarify the practical limitations of IA-DTB chemoproteomics coverage. We observe systematic underrepresentation of specific protein classes and experimental contexts, including secreted proteins and a substantial fraction of insoluble transmembrane proteins that are lost or depleted during standard preparation, and targets limited by low abundance and MS sensitivity. Follow-up experiments underscore both the opportunities and constraints: changing cell lines can recover additional cysteines via altered expression landscapes; fractionation can improve access to membrane/nuclear compartments; and alternative electrophiles can add complementary coverage. Overall, combining all three approaches yielded >30% improvement versus our original protocol in HEK293T alone. However, each comes with added complexity and instrument/sample-prep cost, with diminishing returns for high-throughput screening scenarios. Recent advances have also enabled the proteomic study of secreted proteins (e.g. cytokines)^30^, however require bespoke preparations that are orthogonal to intercellular study. Finally, several limitations of this study primarily reflect the computational approximations required to featurise cysteines at proteome scale. Structural descriptors are derived from static structures and therefore represent only a snapshot of highly dynamic proteins, which is particularly problematic for reactive cysteines often located in flexible functional regions ^31^. These issues may be exacerbated when using predicted structures (e.g., for proteins with low-confidence regions)^18,32,33^, and in turn may propagate error into derived quantities such as p*K*_a_ and other microenvironment features. In addition, cysteine redox state is not explicitly modelled here, and estimates remain sensitive to structural uncertainty and to the limitations of modelling redox biology from static (and predicted) structures alone.

In addition to advancing fundamental knowledge on cysteinome detectability, this work delivers four computational tools to the chemical biology and proteomics community:

1. **ResidueMap** (https://github.com/FrancisCrickInstitute/ResidueMap): A scalable computational pipeline that generates physicochemical features for nucleophilic residues (Cys, Lys, Ser, Tyr) and tryptic peptides across arbitrary proteomes. This tool enables researchers to rapidly extract the chemical and structural descriptors required to build predictive models or prioritise reactive targets across diverse datasets.
2. **esm_embedder** (https://github.com/FrancisCrickInstitute/esm_embedder): A Python program that efficiently generates ESM-2 protein language model embeddings for whole protein sequences, tryptic peptides, individual residue types, and their combinations. It streamlines the integration of rich protein language representations into downstream machine learning workflows without requiring complex setup.
3. **Cysteine detectability predictor** (https://github.com/FrancisCrickInstitute/cysteine-detectability-predictor): A user-friendly application leveraging our pre-trained models to forecast cysteine detectability directly from one or more input protein sequences. By outputting a detailed report for each cysteine residue—including its tryptic peptide sequence, predicted IA-DTB detectability, and a confidence score—this tool empowers researchers to rapidly assess target feasibility and optimize experimental design prior to mass spectrometry analysis.
4. **A distance-based filtering methodology:** A dimensionality reduction approach designed to identify confidently detected peptides across experimental replicates and treatment conditions. This workflow provides a robust, data-driven solution for reducing technical noise and improving data harmonization in large-scale chemoproteomic screens.

By bridging computational prediction with practical chemoproteomics, this study shifts workflow optimisation from empirical trial-and-error to a more data-driven, rational design. Ultimately, these mechanistic insights and open-source tools will empower researchers to maximise target coverage and accelerate the discovery of novel covalent probes and therapeutics.

## Materials and Methods

### Chemoproteomics methods

#### General IA-DTB chemoproteomics workflow

Washed and dried cell pellets were thawed on ice. To the cell pellets was added RIPA lysis buffer (0.6 mL per 100 million cells). The cells were then lysed *via* probe sonication using a Branson probe sonicator (20% amplitude, 5x 5 seconds). The resultant lysate was then clarified via filtration using a Stericup Quick Release Vacuum Driven Filtration system (0.22 µM, ThermoScientific, Cat#: 15780319). Cell lysate concentration was then determined using the Pierce BCA Protein Assay Kit (ThermoScientific, Cat#: 23225) and normalised to 2.5 mg/mL. The cell lysate was then treated according to the established IA-DTB protocol^8^ . Samples were loaded onto EvoTips (5-7.5% loading) and eluted using an Evosep One with a 60 SPD method. Peptides were analysed *via* LC-MS/MS on a Bruker TIMS-ToF Pro2 using a fixed window, DIA method.

### Spectral library generation

Off-line high pH fractionation was performed on pooled DMSO samples for each cell line (HEK293T, Jurkat, eHAP), with the fractions subsequently analysed via 30SPD DDA-PASEF (HEK293T and Jurkat fractions were the ones used in the study by Biggs and Cawood et al^8^). A hybrid library was generated on these combined datasets using the Pulsar search algorithm within Spectronaut. This hybrid library consisted of 122,257 peptide-spectral matches for 68,172 cys-containing peptides on 11,394 proteins. The 16 DIA-PASEF mass spectrometry files generated from HEK293T lysate were then searched against this hybrid library to identify the cysteineome coverage. For reference, the search was performed against the SwissProt human proteome (accessed Q3 2022) which at that time contained 20,577 human proteins, excluding isoforms and retaining homologues.

### Subcellular Fractionation

A washed and dried HEK293T cell pellet (100,000,000 cells) was thawed on ice. Subcellular fractionation was performed using the Subcellular Protein Fractionation Kit for Cultured Cells (ThermoScientific, Cat#: 78840) according to the manufacturers protocol. Subcellular fractions’ lysate concentration was then determined using the Pierce BCA Protein Assay Kit (ThermoScientific, Cat#: 23225) and normalised to 1.0 mg/mL (Cytoplasmic, Membrane and Nuclear fractions) or 0.5 mg/mL (Chromatin fraction). Four aliquots of each fraction (198 µL) were then treated according to the established IA-DTB protocol^8^ . Global proteomics samples were collected from the non-enriched eluent following Neutravidin enrichment. An aliquot of this enrichment (10 µL) was diluted with 0.1% Formic Acid_(aq)_ (90 µL) and loaded directly onto EvoTips (EvoSep, Cat#: EV2018). Prepared samples were loaded onto EvoTips (5-7.5% loading) and eluted using an Evosep One with either a 30 SPD (DDA) or 60 SPD (DIA) method. Peptides were analysed *via* LC-MS/MS on a Bruker TIMS-ToF Pro2 using a fixed window, DIA method. A hybrid library was generated using all DDA and DIA samples across the fractionated HEK293T lysate. This library was generated in Spectronaut using the search algorithm Pulsar. Peptide lengths of 7–52 amino acids and with up to two miscleavages were permitted. The following variable modifications were applied: oxidation (methionine, +15.99 Da), IA-DTB (cysteine, +296.18 Da), acetylation (N-terminus, +42.01 Da), and carbamidomethylation (cysteine, +57.02 Da). All DIA samples for both sub-cellular fractionated and whole lysate control were searched in Spectronaut (v19.1.240806.62635) using this generated hybrid library, following the search settings reported previously^8^. Quality of the cellular fractionation was assessed by measuring abundance of reference proteins in each fraction, as indicated by the kit manufacturer (Figure S11).

### Additional Cell Lines

Eight cell lines (A549, HEK293T, HeLa, hTERT, MCF7, Jurkat, U2OS and U87MG) were cultured, and washed and dried pellets (100,000,000 cells) were collected and stored on ice. To the cell pellets was added RIPA lysis buffer (3 mL per pellet). The cells were then lysed *via* probe sonication using a Branson probe sonicator (20% amplitude, 5x 5 seconds). Cell lysate concentration was then determined using the Pierce BCA Protein Assay Kit (ThermoScientific, Cat#: 23225) and normalised to 1.0 mg/mL. The cell lysate was then treated according to the established IA-DTB protocol^8^. Global proteomics samples were collected from the non-enriched eluent following Neutravidin enrichment. An aliquot of this enrichment (10 µL) was diluted with 0.1% Formic Acid_(aq)_ (90 µL) and loaded directly onto EvoTips (EvoSep, Cat#: EV2018). Prepared samples were loaded onto EvoTips (5-7.5% loading) and eluted using an Evosep One with either a 30 SPD (DDA) or 60 SPD (DIA) method. Peptides were analysed *via* LC-MS/MS on a Bruker TIMS-ToF Pro2 using a fixed window, DIA method. A hybrid library was generated using all DDA and DIA samples across all the cell lines. This library was generated in Spectronaut using the search algorithm Pulsar. Peptide lengths of 7–52 amino acids and with up to two miscleavages were permitted. The following variable modifications were applied: oxidation (methionine, +15.99 Da), IA-DTB (cysteine, +296.18 Da), acetylation (N-terminus, +42.01 Da), and carbamidomethylation (cysteine, +57.02 Da). All DIA samples for all cell lines were searched in Spectronaut (v19.1.240806.62635) using this generated hybrid library, following the search settings reported previously^8^.

### Alternative ABPP Probes

A washed and dried HEK293T cell pellet (500,000,000 cells) was thawed on ice. To the cell pellet was added RIPA lysis buffer (3 mL). The cells were then lysed *via* probe sonication using a Branson probe sonicator (20% amplitude, 5x 5 seconds). The resultant lysate was then clarified via filtration using a Stericup Quick Release Vacuum Driven Filtration system (0.22 µM, ThermoScientific, Cat#: 15780319). Cell lysate concentration was then determined using the Pierce BCA Protein Assay Kit (ThermoScientific, Cat#: 23225) and normalised to 1.0 mg/mL. For each ABPP probe, a series of stock solutions (1, 10 and 50 mM) in DMSO were prepared, and these stocks were aliquoted (2 µL) onto 96-well plates. To the probes was added an aliquot of the prepared HEK293T lysate (198 µL), with the resultant treatments incubated at room temperature for one hour. The treated cell lysates were then processed according to the established IA-DTB protocol^8^ . Prepared samples were loaded onto EvoTips (5-7.5% loading) and eluted using an Evosep One with either a 30 SPD (DDA) or 60 SPD (DIA) method. The DDA samples generated were used to generate optimised DIA-PASEF windows for each ABPP probe, accounting for shifted *m*/*z* and ion mobility characteristics between probes. Window optimisation was performed using the pyDIAD python package and methods reported previously^34^. FragPipe MSFragger^35,36^ open search with PTM-Shepherd^37,38^ was performed on the DDA samples to identify the amino acid selectivity and mass of modification of each probe (Figure S12a). Peptides were then analysed *via* LC-MS/MS on a Bruker TIMS-ToF Pro2 using the optimised DIA method for each probe. A hybrid library for each ABPP probe was generated using all DDA and DIA samples for the respective probe treated lysate. These libraries were generated in Spectronaut using the search algorithm Pulsar. Peptide lengths of 7–52 amino acids and with up to two miscleavages were permitted. The following variable modifications were applied: oxidation (methionine, +15.99 Da), DTB-probe (cysteine, variable mass of modification, Figure S12b), acetylation (N-terminus, +42.01 Da), and carbamidomethylation (cysteine, +57.02 Da). All DIA samples were searched in Spectronaut (v16.0.220606.53000) using their respective hybrid libraries, following the search settings reported previously^8^.

### Computational methods

#### Data preparation

A reference human proteome of 20577 canonical and reviewed human protein sequences was obtained from the Swiss-Prot fraction of UniProt^17^. The reference proteome was identical to the proteome used for chemoproteomics analysis. Each sequence was mapped to its respective AlphaFold2^18^ predicted structural model through the AlphaFold Database^19^. Sequences and structural models were used to calculate features as described below. Data collection from both resources was performed using their public API.

### Labelling of robustly detected peptides

Per-peptide metrics were computed across DMSO replicates: mean signal (Intensity), data completeness (DC; fraction of replicates with non-missing signal) and coefficient of variation across valid replicates (CV). Intensity was log2-transformed and Intensity, DC and CV were rescaled to [0,1] within each MS experiment. The normalised metric vector *m = (I_norm_, DC, CV_norm_)* was then compared to the ideal peptide *(1,1,1)* by calculating their Euclidean distance in 3D space *d* = 0(1 − *I_norm_*)^2^ + (1 − *DC*)^2^ + (1 − *CV_norm_*)^2^. A bimodal Gaussian mixture model was fitted to the empirical distribution of distances, and the intersection of the two Gaussian densities was taken as the robustness cutoff *d_cut_*, and peptides with *d ≤ d_cut_* were labelled robustly detected. With a careful look at the data distribution in 3D space (Figure 2a, left), one can argue that there are still several confidently detected peptides of low intensity that are left out of the spherical inclusion zone. The method can be applied at the peptide level or the cysteine site level, provided that all the metrics of all tryptic peptides carrying the same cysteines, are averaged out. For cysteine-site reporting, peptide metrics were averaged across all tryptic peptides mapping to the same cysteine prior to normalisation and classification. The algorithm requires at least four DMSO replicates to stabilise DC/CV estimates. Diagnostic plots (3D metric scatter, distance histogram with GMM fit and intersection) were generated for quality check (Figure 2a,b). Definitions used in validation: legacy filters = Intensity ≥100, DC ≥0.75, CV ≤0.40; robustly liganded hits = robustly detected sites with ≥2-fold mean signal loss (log2 fold change ≥1) and −log10(p) ≥1.3.

### Calculation of features

Below is a breakdown of each feature type calculated by our *ResidueMap* peptide featurisation pipeline (https://github.com/FrancisCrickInstitute/ResidueMap) and the respective calculation methods:

• Cysteine conservation (‘conservation’): PSI-BLAST was used to construct a Position-Specific Scoring Matrix (PSSM) for a given sequence, using all Swiss-Prot protein sequences as a reference database, with 2 search iterations. Conservation scores were then extracted from the matrix.

• Secondary structure propensities (‘pred_helix’, ‘pred_strand’, ‘pred_turn’, ‘pred_coil’) and predicted solvent accessibility (‘pred_sasa’): The same PSSM was used as input to the program SPINE X^39^ to predict secondary structure propensities (probabilities of observing an α-helix, β-strand, turn or coil) and Solvent Accessible Surface Area (pSASA) via deep learning.

• Explicit solvent accessibility (‘sasa’): The Shrake and Rupley algorithm with a standard probe radius of 1.4Å was used to calculate the Solvent Accessible Surface Area from structure.

• Predicted Part-Sphere Exposure (‘ppse’): pPSE values for cysteines were obtained from published data^9^.

• Cysteine pKa (‘pka’) and disulfide flag (‘in_disulfide’): The program PROPKA3^20^ was used to compute residue pKa values, using structures as input. A NaN value was assigned to cysteines engaged in disulfide bonds.

• Cysteine environment descriptors (‘rA_nres_all’, ‘rA_nres_reactive’, ‘rA_nres_non_reactive’, ‘rA_mean_pka’, ‘rA_mean_sasa’, ‘rA_total_sasa’ where *r* is radius): Descriptors of neighbouring residues in a spherical area around a cysteine residue (for radii *r* of 4, 6, 8 and 10Å) were calculated by applying a k-d tree neighbour search algorithm implemented in the Biopython package, in combination with pKa and SASA calculations as described above. As ‘reactive’ and ‘non-reactive’ residues we define [Cys, Asp, Glu, His, Lys, Arg, Ser, Thr, Tyr] and [Ala, Asn, Gly, Ile, Leu, Met, Phe, Pro, Trp, Val, Gln] respectively.

• Pocket descriptors (‘in_pocket’, ‘pocket_score’, ‘pocket_druggability’, ‘pocket_sasa’, ‘pocket_volume’, ‘pocket_hydrophobicity’, ‘pocket_charge’): The program Fpocket^40^ was applied to identify ligandable pockets in each protein structure. Fpocket identifies surface cavities on proteins by rolling spherical probes (alpha spheres) of varying radii within a defined range. An alpha sphere is placed whenever it simultaneously touches four protein atoms. By default, a pocket is considered valid if it contains at least 35 alpha spheres. In our analysis, the probe radii were set between 3.0 and 5.0 Å. A cysteine residue was classified as residing within a pocket (in_pocket=1) if the sulfur atom of its thiol group was located within 1.5 Å of the nearest alpha sphere. From all pockets satisfying this constraint, we selected the one whose edge is closest to the thiol as the residing pocket. Fpocket parameters were determined empirically, based on inspection of multiple examples of cysteine-binding events.

• Protein expression levels (‘mean_expression’, ‘median_expression’, ‘HEK-293_expression’, ‘Jurkat_expression’): MS-1 signal was used as a measure of protein abundance from global proteomics experiments deposited in ProteomicsDB^16^. Expression levels across multiple commonly used cell lines and multiple experiments were collected from ProteomicsDB and the mean and median expression were calculated. Expression levels for HEK-293T and Jurkat were also included in the feature set.

• Tryptic peptide descriptors: The human proteome was *in silico* digested using the trypsin KR|P rule (cleavage after R or K, but not before P) allowing up to two missed cleavages. For each tryptic peptide, three types of descriptors were calculated:

`o` Basic descriptors (‘pep_seq’, ‘pep_length’, ‘pep_n_miscleavages’): Peptide sequence, length and number of missed cleavags from *in silico* tryptic digestion.
`o` Residue type descriptors (’pep_n_hydrophobic’, ’pep_n_hydrophilic’, ’pep_n_positive’, ’pep_n_negative’, ’pep_n_aromatic’, ’pep_n_his’, ’pep_n_pro’, ’pep_n_cys’, ’pep_f_hydrophobic’, ’pep_f_hydrophilic’, ’pep_f_positive’, ’pep_f_negative’, ’pep_f_aromatic’, ’pep_f_his’, ’pep_f_pro’, ’pep_f_cys’): Counts and frequencies of different residue chemistries and types in a peptide.
`o` Chemical descriptors (’pep_aliphatic_index’, ’pep_boman_index’, ’pep_charge’, ’pep_hydrophobicity’, ’pep_instability_index’, ’pep_pI’, ’pep_unmodified_mass’, ’pep_modified_mass’, ’pep_MolLogP’, ’pep_MolMR’, ’pep_TPSA’, ’pep_FractionCSP3’): Physicochemical properties of peptides calculated by rdkit^41^ and the *peptides*python package (https://github.com/althonos/peptides.py). As ‘pep_modified_mass’ and ‘pep_unmodified_mass’ we define the mass of the peptide with and without including oxidised Met, DTB-modified Cys and acetylated N-termini modifications. ‘pep_MolLogP’ and ‘pep_MolMR’ are the log-transformed partition coefficient, a measure of lipophilicity, and molar refractivity, a measure of polarisability, both calculated using Crippen’s method^42^ in *rdkit*. ‘pep_TPSA’ and ‘pep_FractionCSP3’ are the topological polar surface area of the peptide and its fraction of sp^3^ hybridised carbon atoms, again calculated with *rdkit*.

• ESM-2 protein sequence embedding: All Protein Language Models embeddings were derived using Facebook’s esm2_t36_3B_UR50D model^43^. Sequence embeddings were obtained by averaging all token values. Individual cysteine embeddings were similarly obtained by taking the corresponding token embedding after the last transformer block. ESM embeddings are produced at three levels for each cysteine. First mean embedding is computed for the whole protein the cysteine is part of, averaging over all amino acids in the protein to obtain a 2560-sized vector per protein. Then the individual token embedding corresponding to the target cysteine is extracted as well leading again to a 2560-sized vector. Finally, the mean embedding of the peptide (shortest tryptic peptide or average of all tryptic peptides for a cysteine) in which the embedding is computed by averaging all amino acids present in the peptide sequence leading to the final 2560-sized vector. All three embeddings are concatenated or averaged to represent a cysteine and an associated peptide. Embeddings were computed using GPU parallelization on an HPC cluster. Sequences longer than 11000 amino acids were discarded but no sequences were cropped.

The ‘all features’ set includes proteomic abundance data, structural information and all peptide features. ‘Peptide only’ and ‘Structural only’ contain, respectively, peptide features and structural features, in addition to expression data. Curated features consist in the following 17 features ’pep_length’, ’pep_modified_mass’, ’pep_pI’, ’pep_MolLogP’, ’mean_expression’, ’conservation’, ’plddt’, ’pka’, ’ppse’, ’sasa’, ’6A_total_pka’, ’pep_n_cys’, ’pep_n_pro’, ’pep_n_positive’, ’pep_n_negative’, ’pep_n_aromatic’, ’pep_n_miscleavages’. All feature pairs are built by forming pairs of combinations without repeat of the curated features. ‘ESM’ features are the concatenated representations of the protein, cysteine and peptide as detailed in the featurisation pipeline with the exclusion of any additional features. The featurisation pipeline was deployed on HPC for efficient parallelisation over the whole proteome.

### ML training/validation data preparation

The train-test split in each of the 5-fold cross validation was set to 80/20 percent. Balanced datasets were always used both in detectability and ligandability by downsampling the majority class to match the number of samples in the minority class. The test/train sets are built so that one ‘uniprot_id’ appears only in either the training or test dataset, but never both. This is to minimise the data leakage that would occur from having the same protein in both sets. The detectability dataset was built by taking the 25,494 peptides robustly detected via IA-DTB as positives. Negative samples were sampled without replacement from the 1,170,174 undetected peptides, among all 1,192,668 peptides derived from *in silico* tryptic digestion (see further above).

### Training/testing set sampling and feature filtering strategy

The featurised cysteinome data was split into test and train sets by retaining the same number of negatives as the number of positives (balanced dataset). Random sampling resulted in a heavily unbalanced dataset with negatives dominating, which resulted in a model biased towards negative answers, therefore we decided to proceed with a balanced set for training, and for testing, we used both a balanced set and a random, unbalanced set. In all cases, to prevent data leakage from the training to the test set, we made sure no peptides from the same protein are included in both splits. Only peptides with no missed cleavages were used, as the number of detected peptides with missed cleavages is vanishingly small compared to the virtual space of possible missed cleavages, which over-skews the dataset.

### ML model implementation

Five initial models were tested (see Table 1) for selection of the most optimal: linear regression, logistic regression, random forest, SVM and Gradient Boosted Tree (CatBoost). Those were trained with default hyperparameters, only on HEK293T derived labels, with a balanced training set, and both balanced and unbalanced testing sets. Median imputation of missing values across rows and columns was applied for linear regression and random forest, while Z-score normalisation was applied for logistic regression and SVM. All five models were implemented through the scikit-learn Python package. CatBoost (version 1.2.8) was used for follow-up without hyperparameter tuning. CatBoost natively handles boolean and categorical variables in addition to NaNs which are present in the feature table. Predictions with the ESM-2 embedding features^44^ were performed with an mlp^26^ as implemented by MLPClassifier in scikit-learn (version 1.5.1)^45^ with two hidden layers of 50 neurons each, default hyperparameters and trained using an early stopping criterion on a 10% validation set taken from the training set. Dummy classifiers in the PR and ROC curves were set to output uniform probabilities, regardless of the input, using scikit-learn version 1.5.1. Finally, SHAP plots were obtained using the SHAP package (version 0.46.0) in combination with the ExplainableBoostingClassifier which is tree-based Generalised Additive Model (GAM)^46^ as implemented in the InterpretML (version 0.6.6) package^47^. Hyperparameters were left to default, but the number of interactions was set to 0 to disable feature interactions and facilitate feature importance interpretations. All experiments and data preprocessing were performed in Python 3.10. Beeswarm plots were obtained using the GAM and the SHAP package using the TreeExplainer and the beeswarm function. The code for reproducing the models used in this study is available at: https://github.com/FrancisCrickInstitute/chemoproteomics-interpretability.

## Author contributions

**IGR**: Conceptualisation, Methodology, Formal analysis, Software, Investigation, Validation, Writing – Original Draft preparation; **HW**: Conceptualisation, Methodology, Data curation, Writing – Review & Editing; **AC**: Methodology, Software, Writing – Review & Editing; **HL**: Methodology, Software, Writing – Review & Editing; **JD**: Software; **WMC, GB, EC:** Methodology, Data curation; **LN**: Conceptualisation, Validation; **JL**: Conceptualisation; **PM, MG:** Data curation; **AS, KA, MS**: Resources, Project Administration; **KR, SB, DH, JB**: Conceptualisation, Supervision, Resources, Project Administration, Writing – Review & Editing

## Acknowledgements

This work was supported by the Francis Crick Institute, which receives its core funding from Cancer Research UK (CC2075, CC2000), the UK Medical Research Council (CC2075, CC2000) and the Wellcome Trust (CC2075, CC2000. This project has been funded by a Prosperity Partnership grant from the Engineering and Physical Sciences Research Council (EPSRC), EP/V038028/1. We also thank GSK for its commitment to support fundamental discovery research through the initial establishment of the Crick-GSK LinkLabs partnership and its contribution to the EPSRC Prosperity Partnership grant.

