## Supplementary Information for "Determinants of cysteine coverage by chemoproteomics"

**Running Title**

Cysteine coverage prediction by chemoproteomics

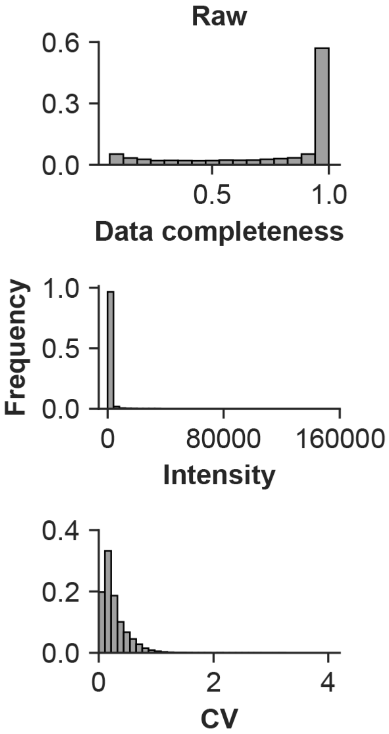


*Figure S1: Distributions of LC-MS quality control metrics for DMSO-treated samples across four replicates, in their raw (left column plots) and normalised/transformed format (right column plots)*

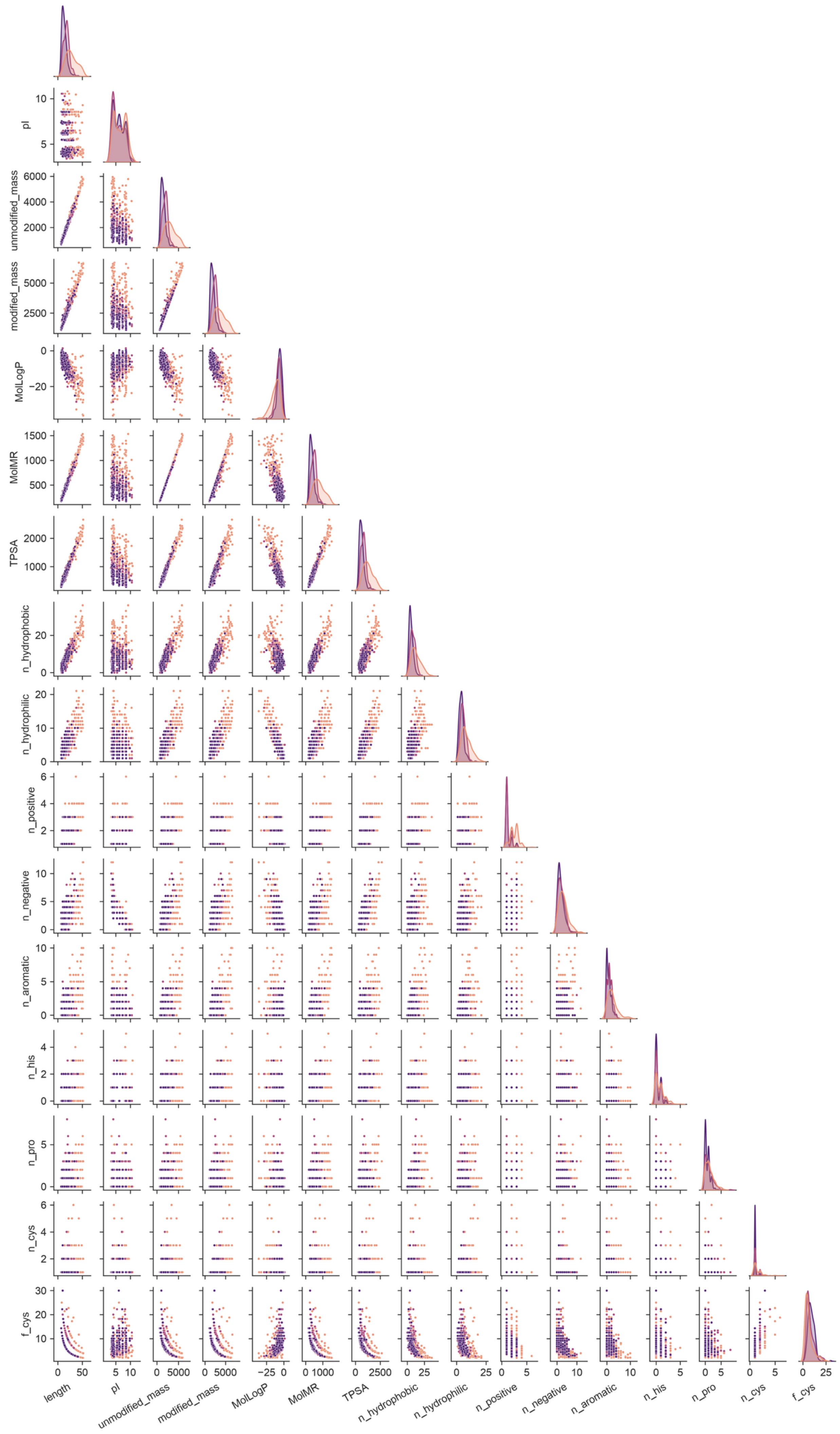
length
pI
unmodified_mass
modified_mass
MolLogP
MolMR
TPSA
n_hydrophobic
n_hydrophilic
n_positive
n_negative
n_aromatic
n_his
n_pro
n_cys
f_cys

*Figure S2: All vs all correlation scatter plots of peptide features, resolved by detectability. Plots on the diagonal show the distributions of each feature.*

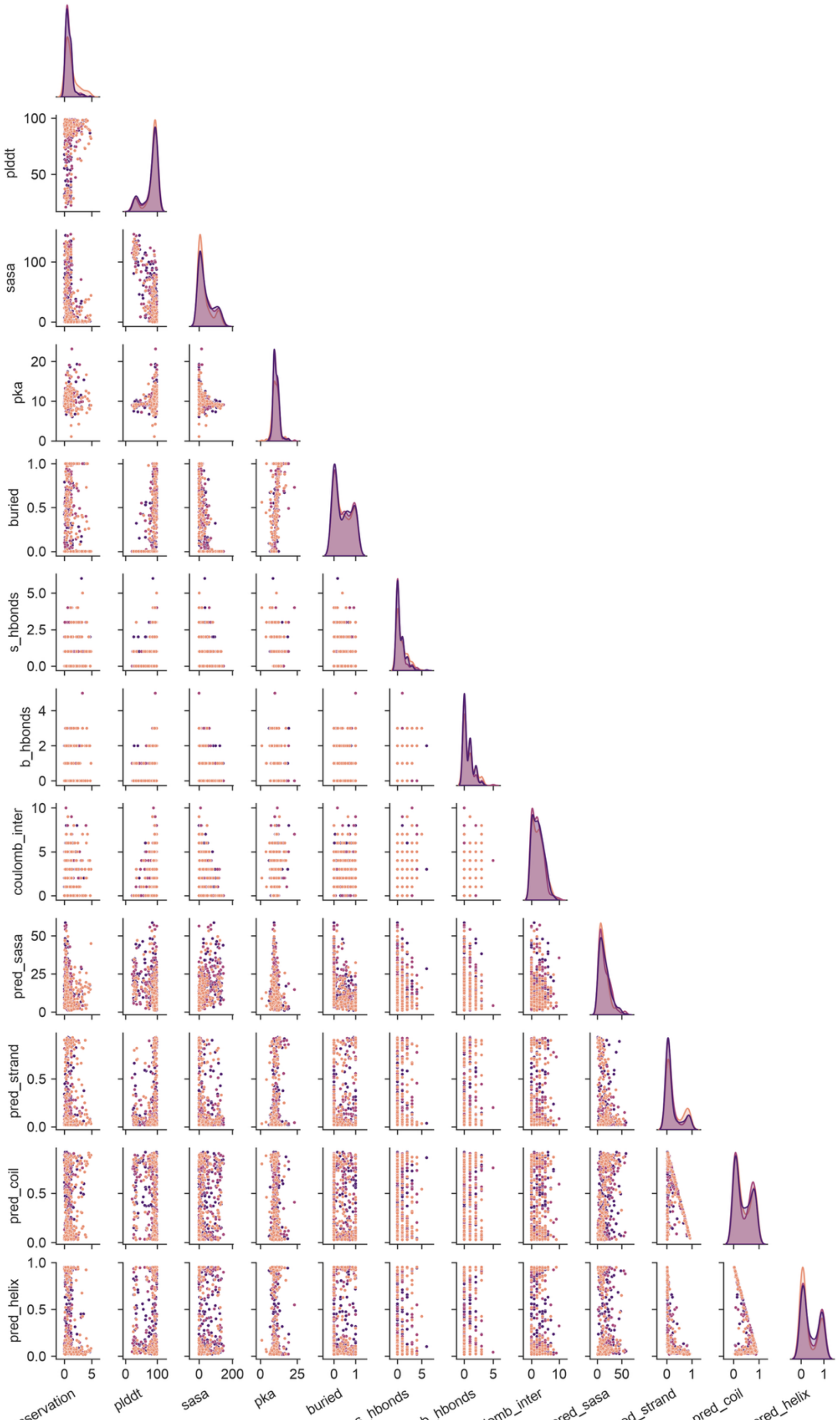


*Figure S3: All vs all correlation scatter plots of cysteine structural features, resolved by detectability. Plots on the diagonal show the distributions of each feature.*

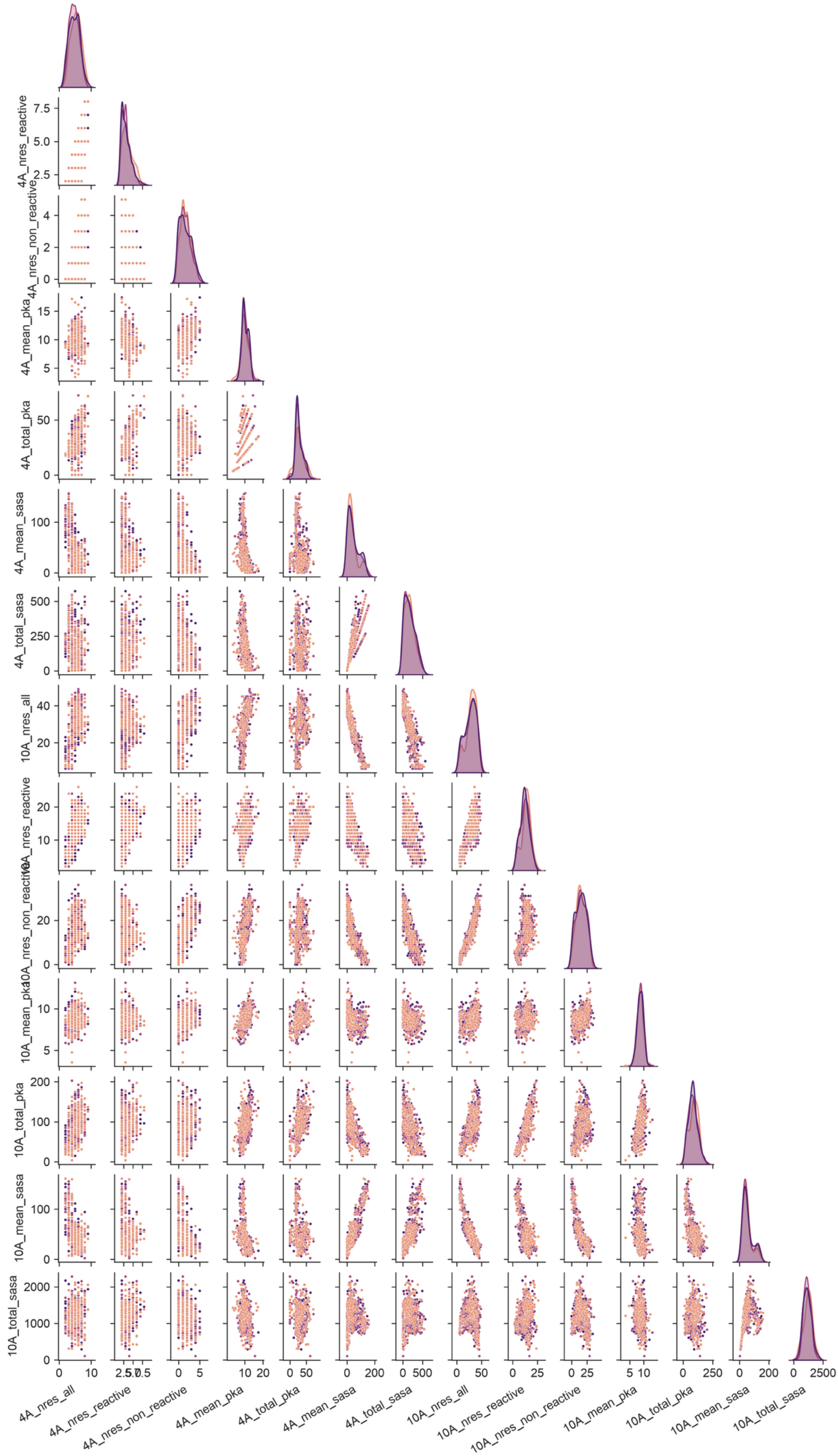

4A_nres_reactive
4A_nres_non_reactive
4A_mean_pka
4A_total_pka
4A_mean_sasa
4A_total_sasa
10A_nres_all
10A_nres_reactive
10A_nres_non_reactive
10A_mean_pka
10A_total_pka
10A_mean_sasa
10A_total_sasa
4A_nres_all
4A_nres_reactive
4A_nres_non_reactive
4A_mean_pka
4A_total_pka
4A_mean_sasa
4A_total_sasa
10A_nres_all
10A_nres_reactive
10A_nres_non_reactive
10A_mean_pka
10A_total_pka
10A_mean_sasa
10A_total_sasa

*Figure S4: All vs all correlation scatter plots of cysteine environment structural features, resolved by detectability. Plots on the diagonal show the distributions of each feature.*

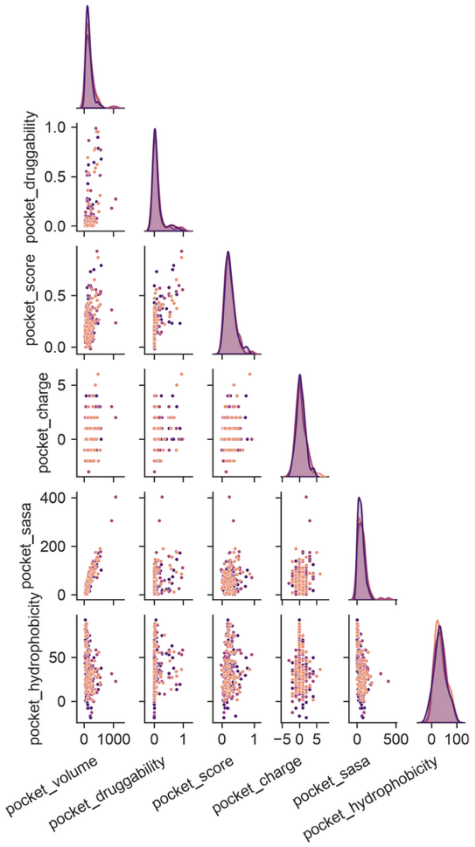


*Figure S5: All vs all correlation scatter plots of cysteine closest pocket features, resolved by detectability. Plots on the diagonal show the distributions of each feature.*

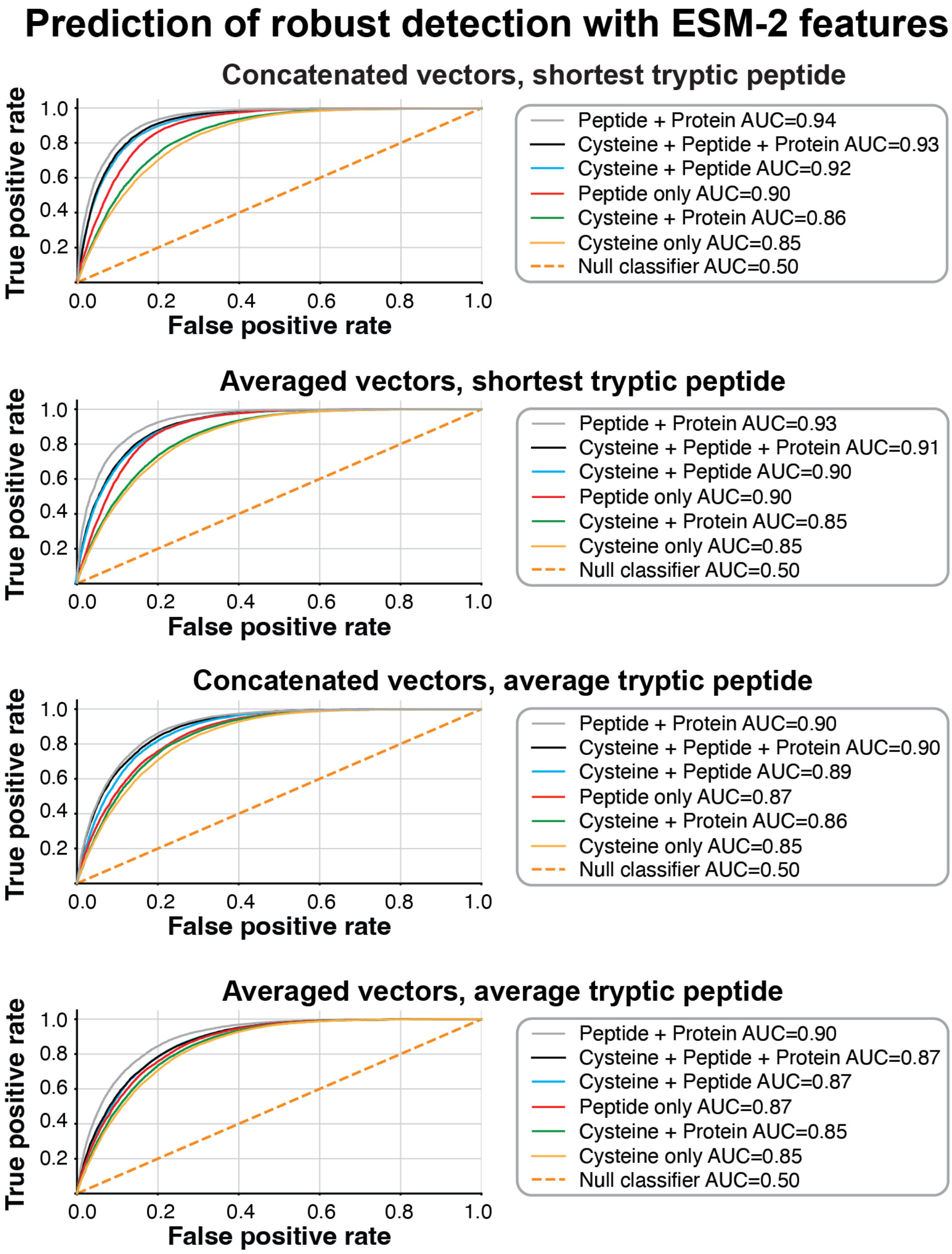


*Figure S6: Benchmark of different embedding combinations, tryptic peptide embedding calculations and vector merging strategies. We found that concatenation of embedding vectors and the use of the shortest tryptic peptide for a cysteine, provide an overall optimal performance.*

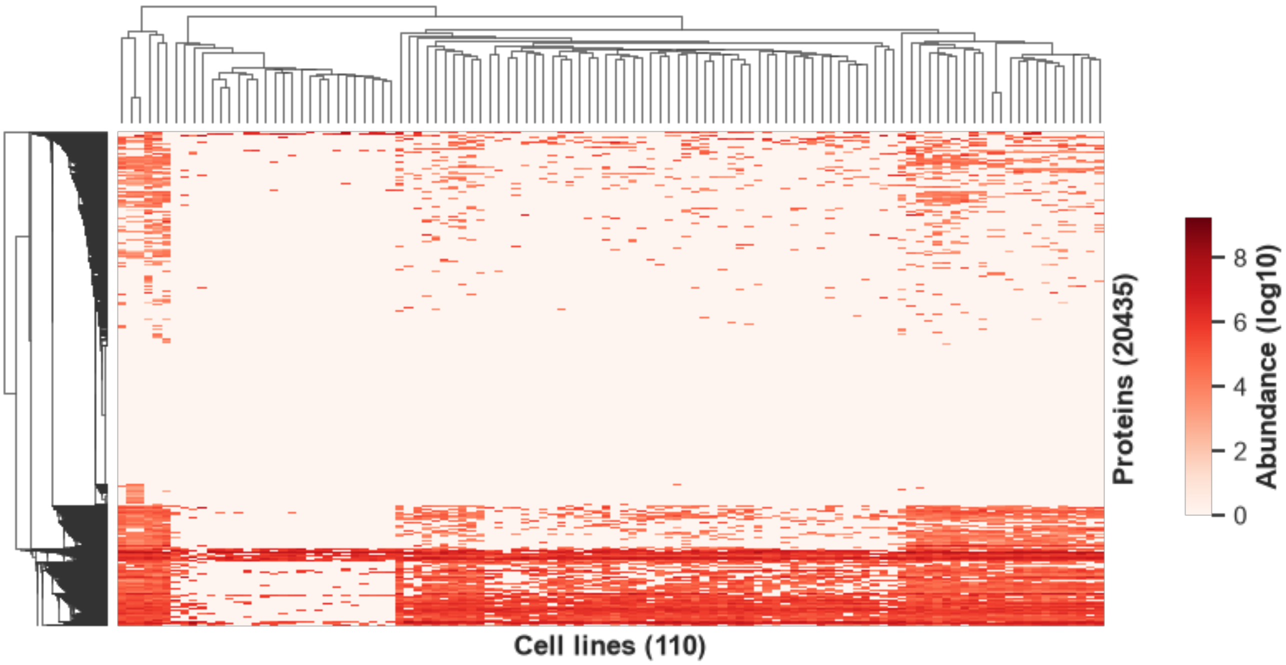


*Figure S7: Heatmap of protein expression (y axis) across cell lines in ProteomicsDB (x axis), highlighting the large fraction of the proteome that is differentially expressed.*

### IA vs IA1 - Peptide Property Comparison

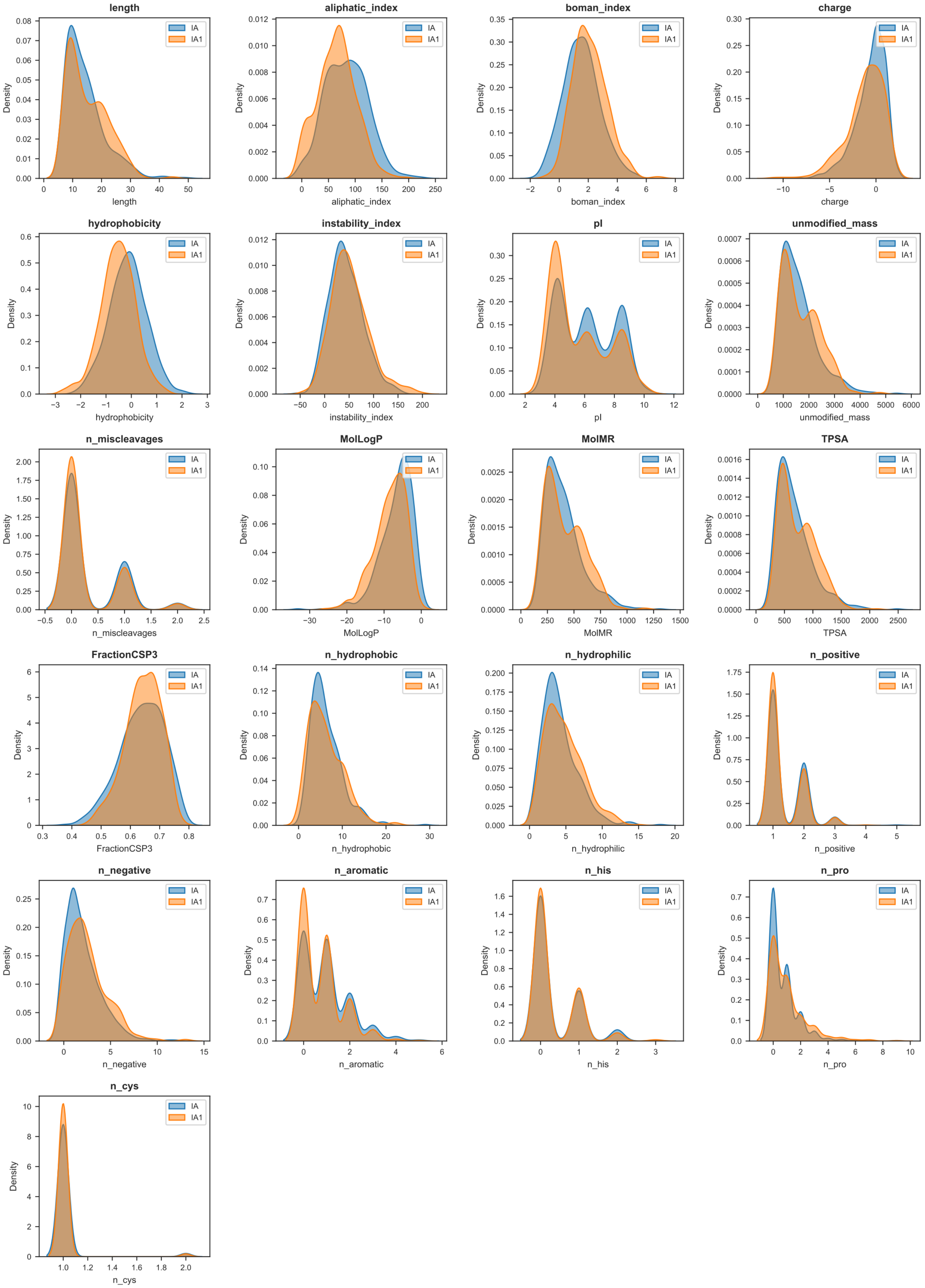

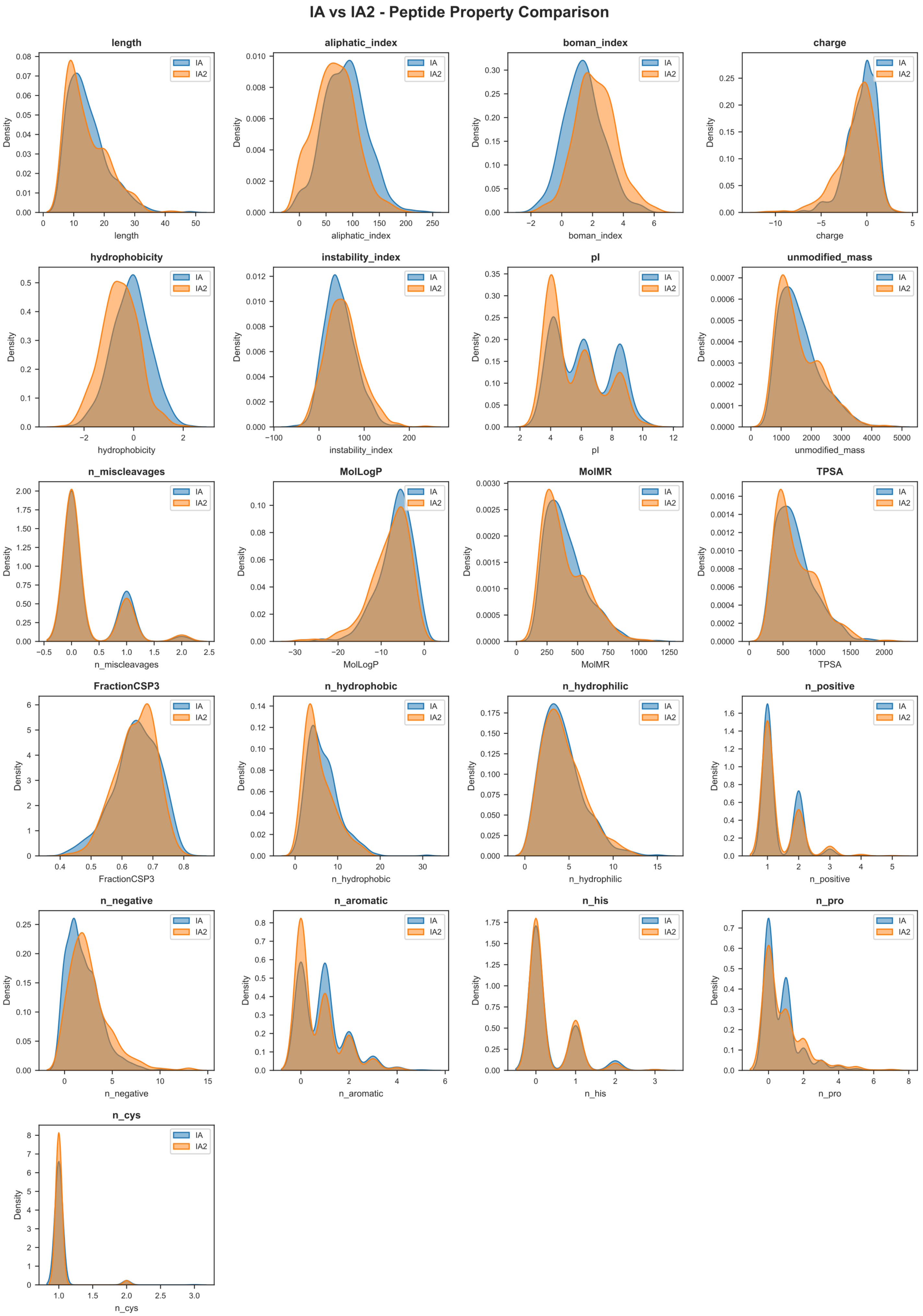
IA vs IA2 - Peptide Property Comparison
length
aliphatic_index
boman_index
charge
hydrophobicity
instability_index
pI
unmodified_mass
n_miscleavages
MolLogP
MolMR
TPSA
FractionCSP3
n_hydrophobic
n_hydrophilic
n_positive
n_negative
n_aromatic
n_his
n_pro
n_cys
Density
IA
IA2

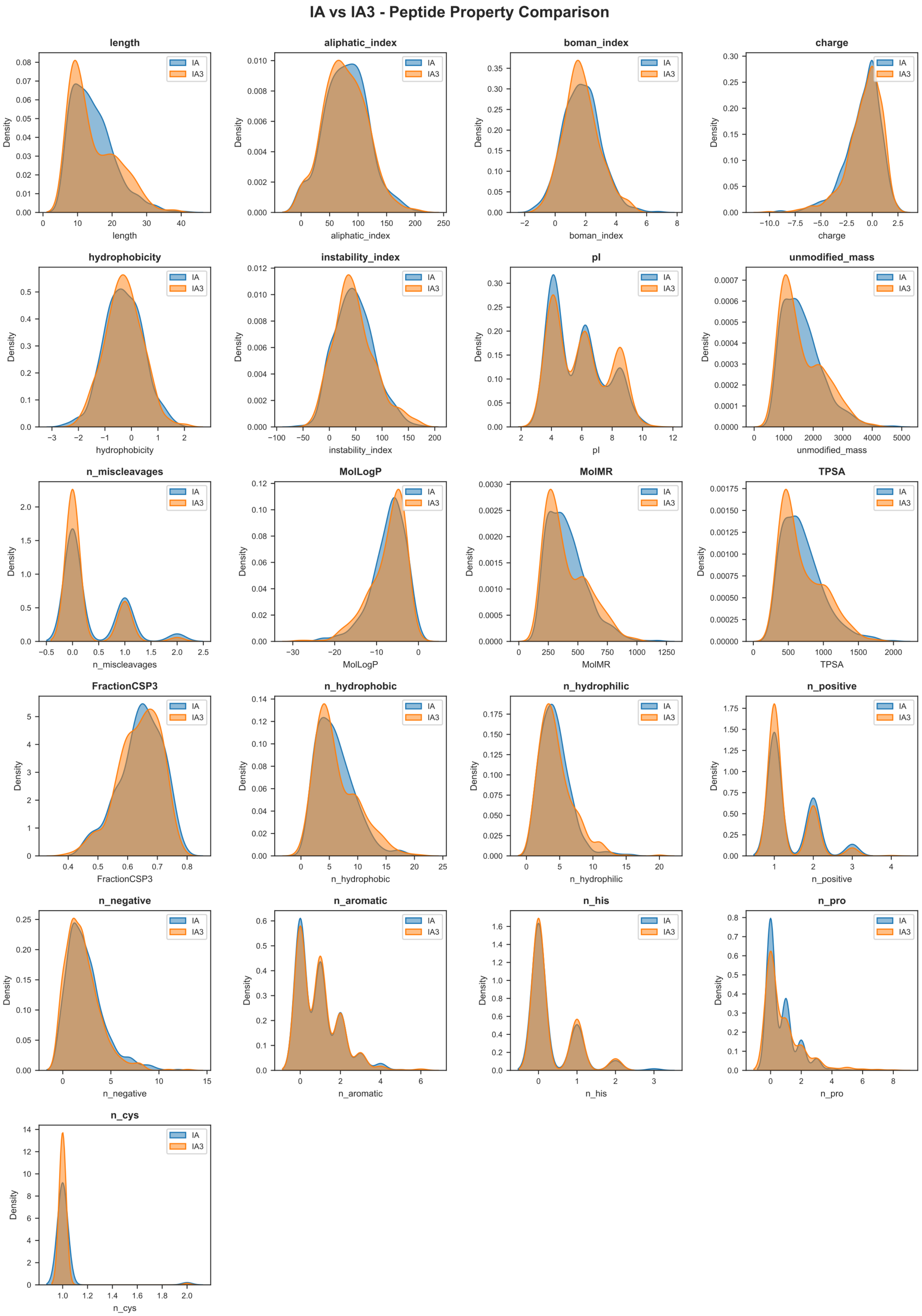
IA vs IA3 - Peptide Property Comparison
length
aliphatic_index
boman_index
charge
hydrophobicity
instability_index
pI
unmodified_mass
n_miscleavages
MolLogP
MolMR
TPSA
FractionCSP3
n_hydrophobic
n_hydrophilic
n_positive
n_negative
n_aromatic
n_his
n_pro
n_cys
Density
IA
IA3

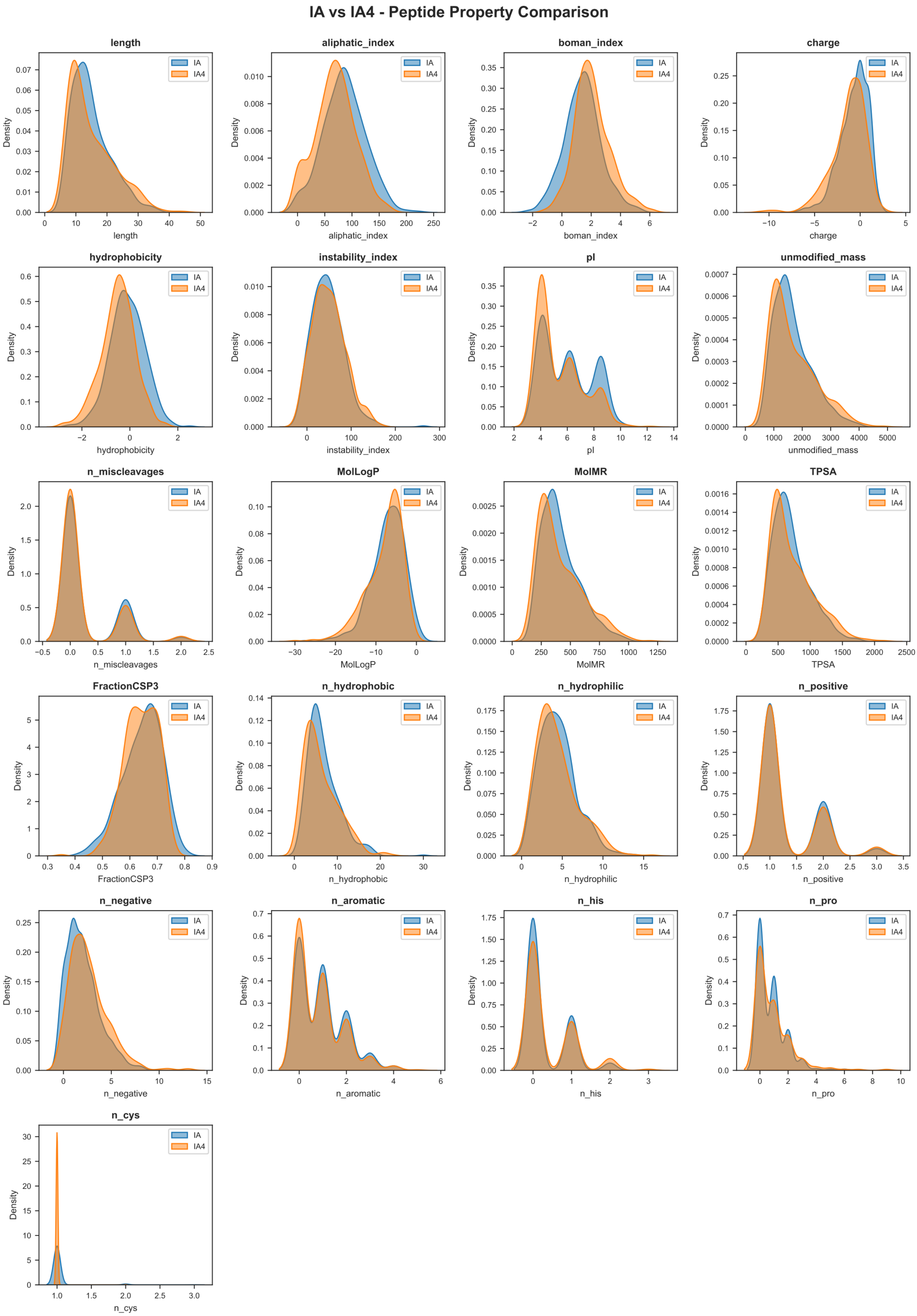
IA vs IA4 - Peptide Property Comparison
length
aliphatic_index
boman_index
charge
hydrophobicity
instability_index
pI
unmodified_mass
n_miscleavages
MolLogP
MolMR
TPSA
FractionCSP3
n_hydrophobic
n_hydrophilic
n_positive
n_negative
n_aromatic
n_his
n_pro
n_cys
Density
IA
IA4

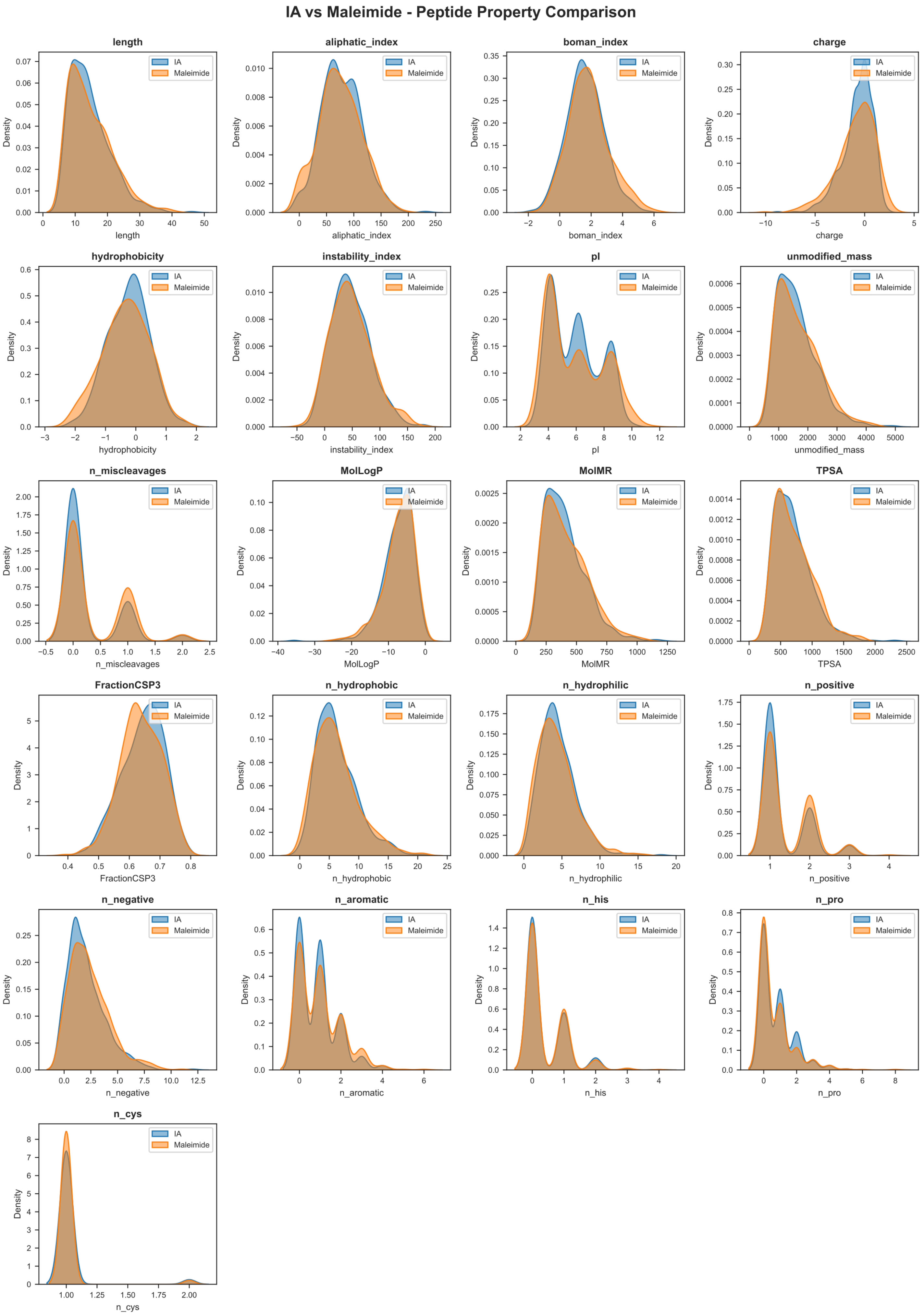
IA vs Maleimide - Peptide Property Comparison
length
aliphatic_index
boman_index
charge
hydrophobicity
instability_index
pI
unmodified_mass
n_miscleavages
MolLogP
MolMR
TPSA
FractionCSP3
n_hydrophobic
n_hydrophilic
n_positive
n_negative
n_aromatic
n_his
n_pro
n_cys
Density
IA
Maleimide

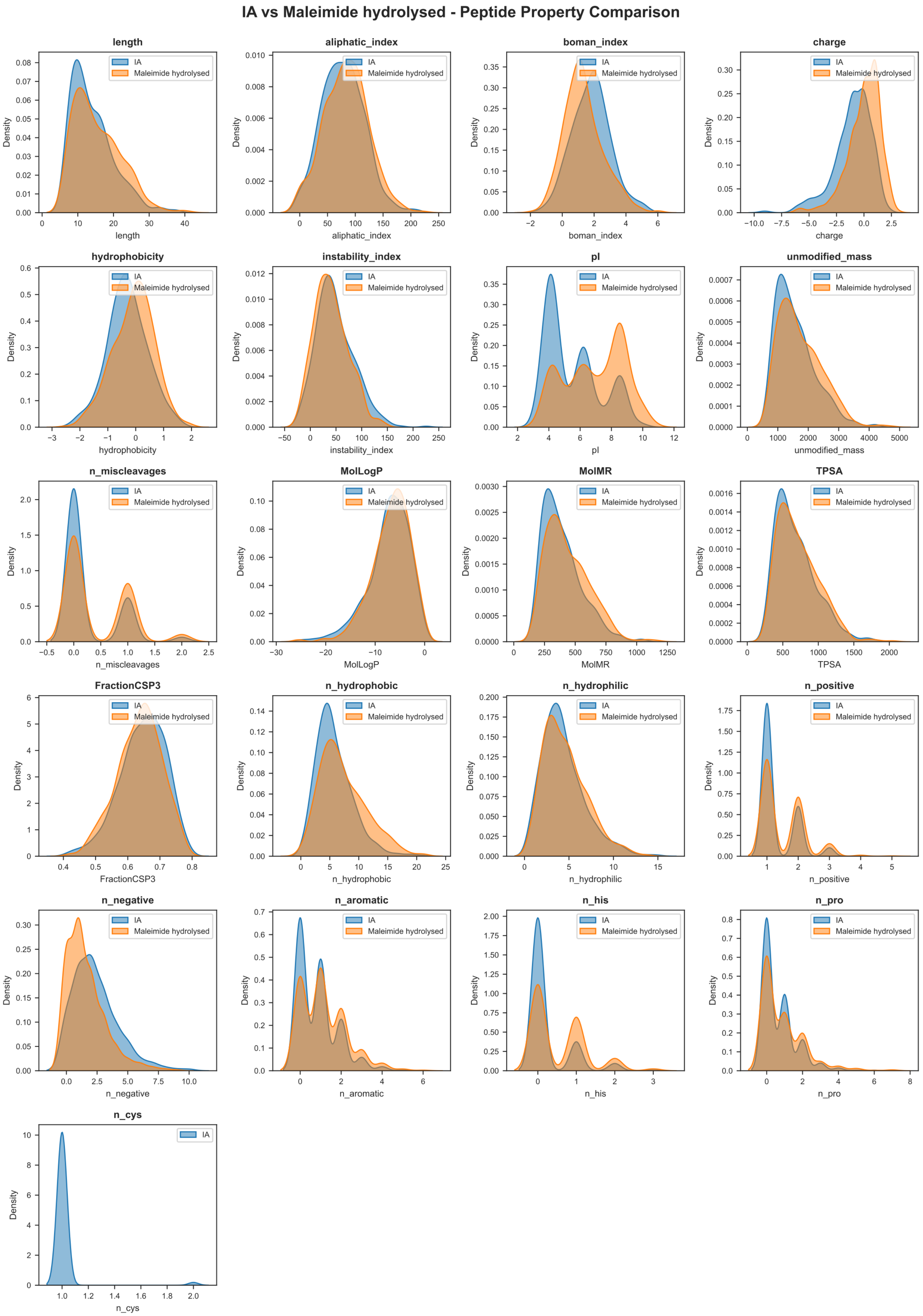
IA vs Maleimide hydrolysed - Peptide Property Comparison
IA
Maleimide hydrolysed
Density
length
aliphatic_index
boman_index
charge
hydrophobicity
instability_index
pI
unmodified_mass
n_miscleavages
MolLogP
MolMR
TPSA
FractionCSP3
n_hydrophobic
n_hydrophilic
n_positive
n_negative
n_aromatic
n_his
n_pro
n_cys

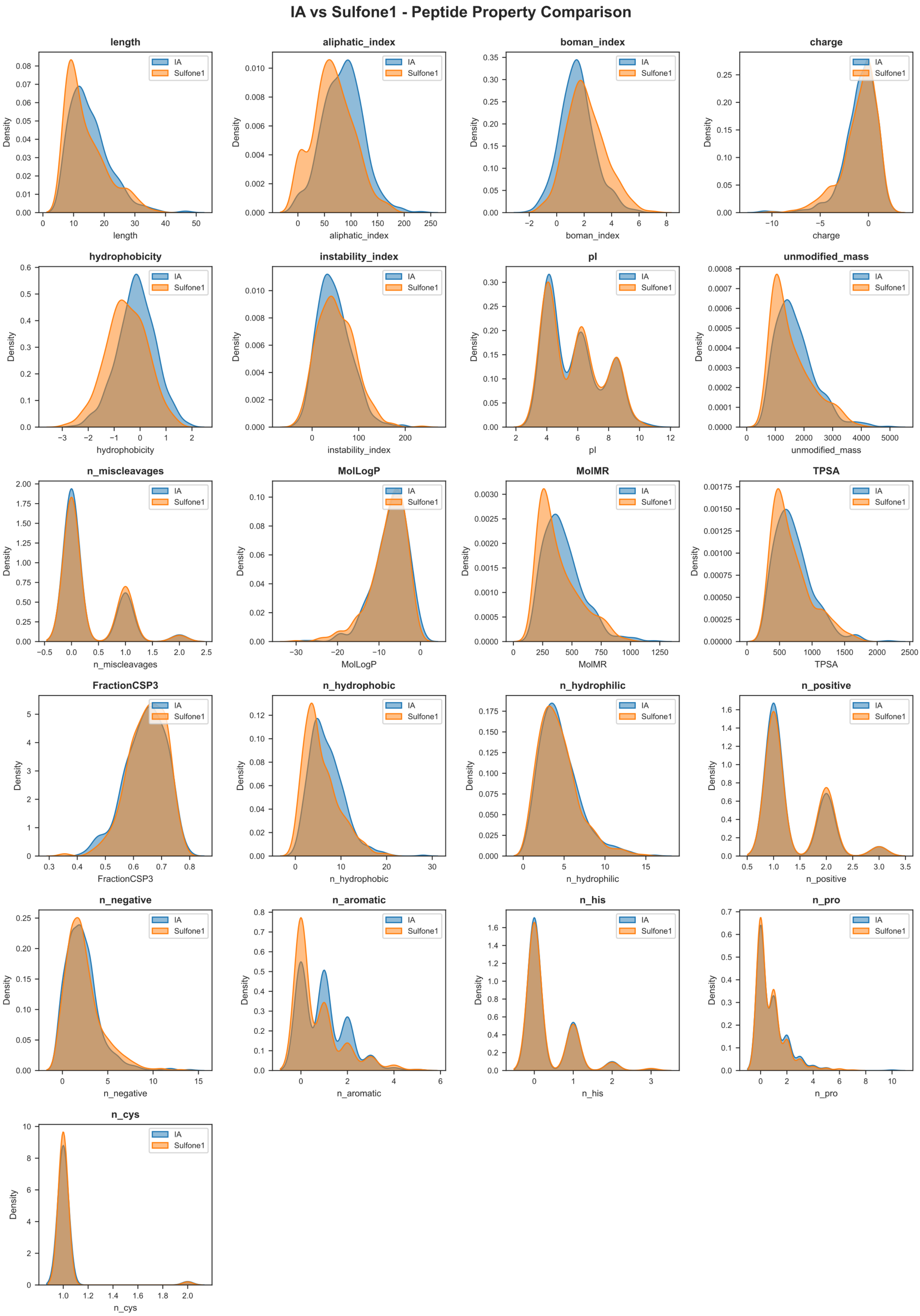
IA vs Sulfone1 - Peptide Property Comparison
IA
Sulfone1
Density
length
aliphatic_index
boman_index
charge
hydrophobicity
instability_index
pI
unmodified_mass
n_miscleavages
MolLogP
MolMR
TPSA
FractionCSP3
n_hydrophobic
n_hydrophilic
n_positive
n_negative
n_aromatic
n_his
n_pro
n_cys

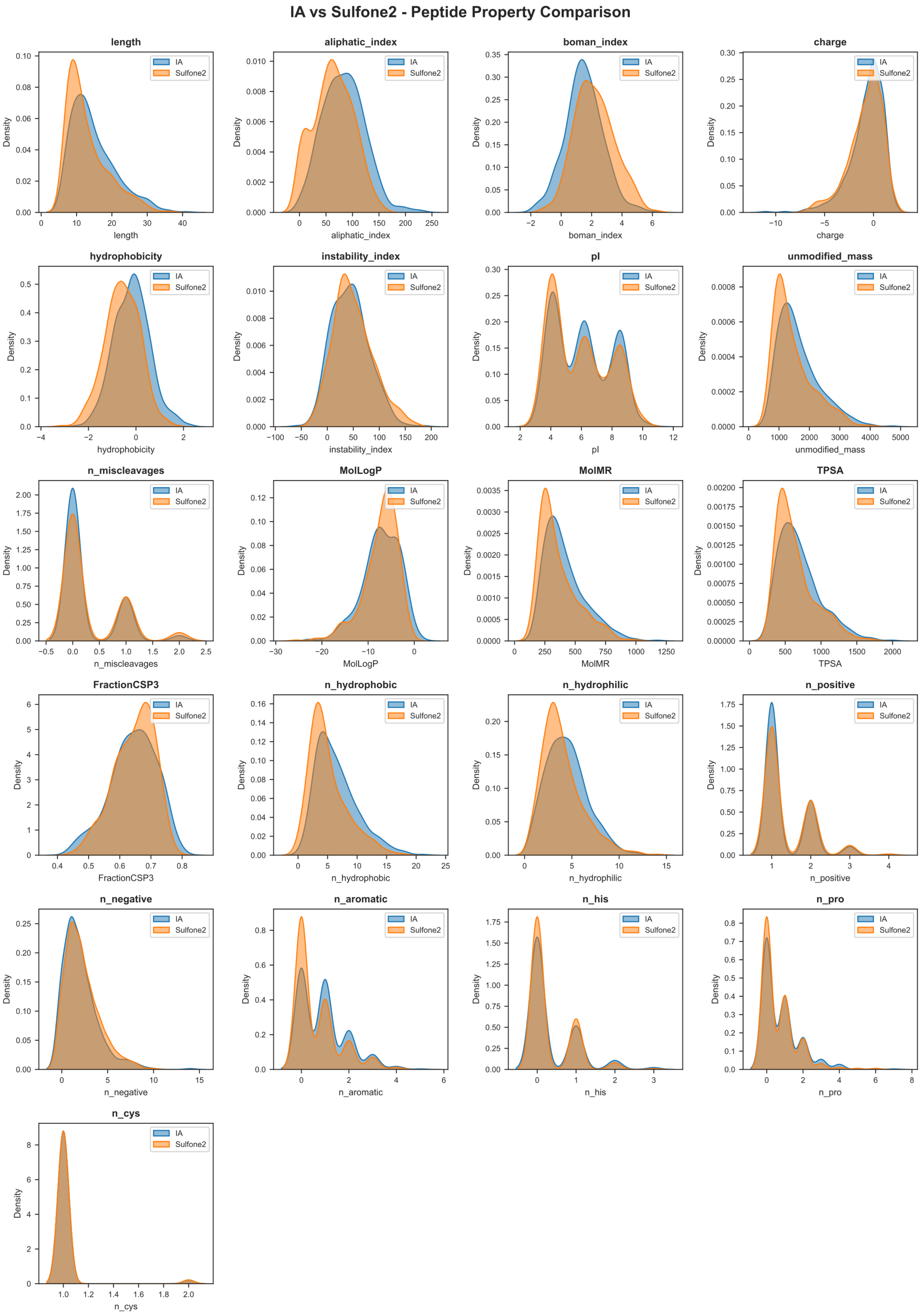
IA vs Sulfone2 - Peptide Property Comparison
length
aliphatic_index
boman_index
charge
hydrophobicity
instability_index
pI
unmodified_mass
n_miscleavages
MolLogP
MolMR
TPSA
FractionCSP3
n_hydrophobic
n_hydrophilic
n_positive
n_negative
n_aromatic
n_his
n_pro
n_cys
Density
IA
Sulfone2

*Figure S8: Peptide feature distributions in 500 randomly selected, uniquely detected peptides by IA-DTB vs IA1-DTB, IA2-DTB, IA3-DTB, IA4-DTB, Maleimide2-DTB, Maleimide2 hydrolysed-DTB, Sulfonamide1-DTB and Sulfonamide2-DTB. Distributions are shown as Kernel Density Estimation plots.*

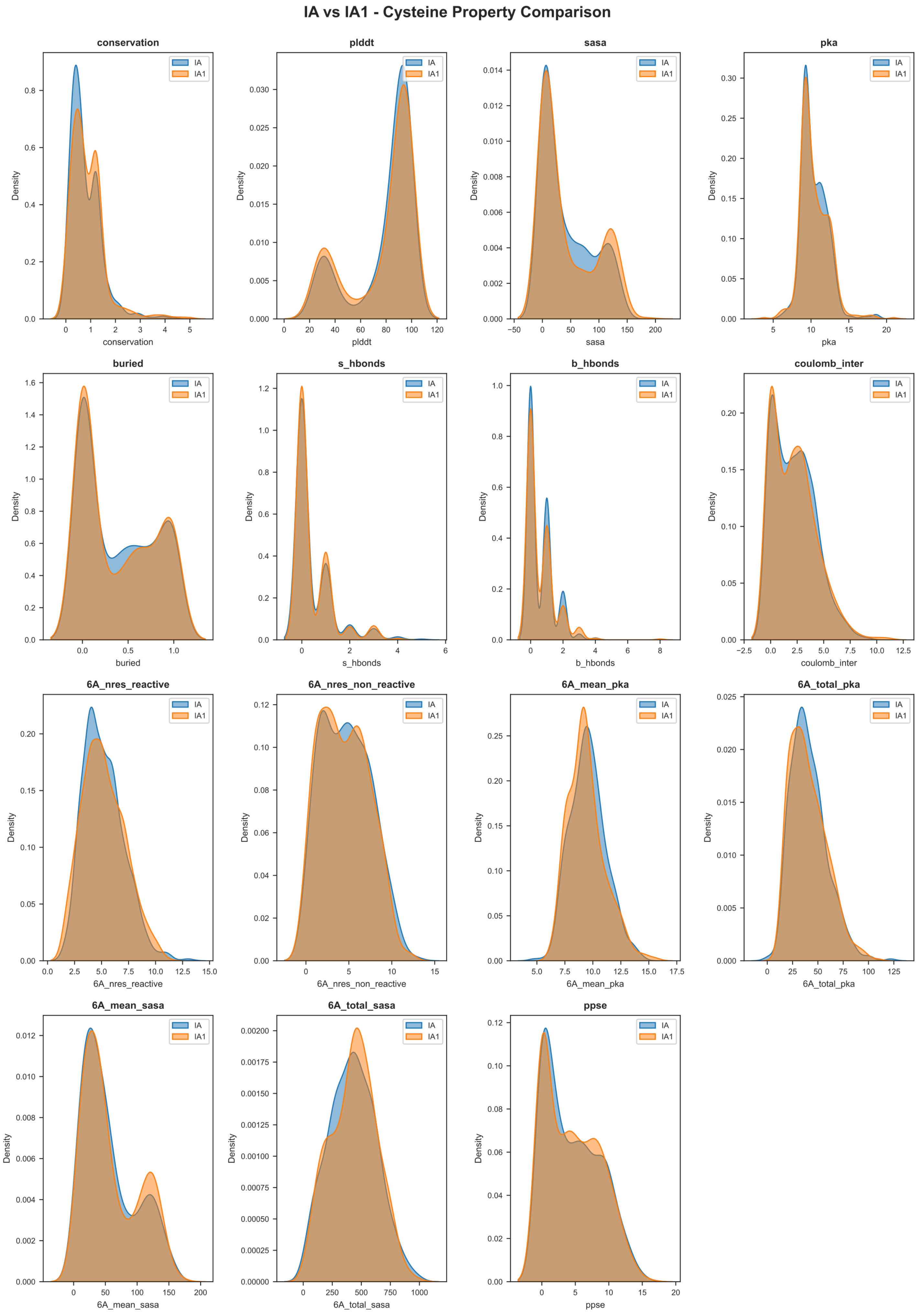
IA vs IA1 - Cysteine Property Comparison
conservation
pldtt
sasa
pka
buried
s_hbonds
b_hbonds
coulomb_inter
6A_nres_reactive
6A_nres_non_reactive
6A_mean_pka
6A_total_pka
6A_mean_sasa
6A_total_sasa
ppse
Density
IA
IA1

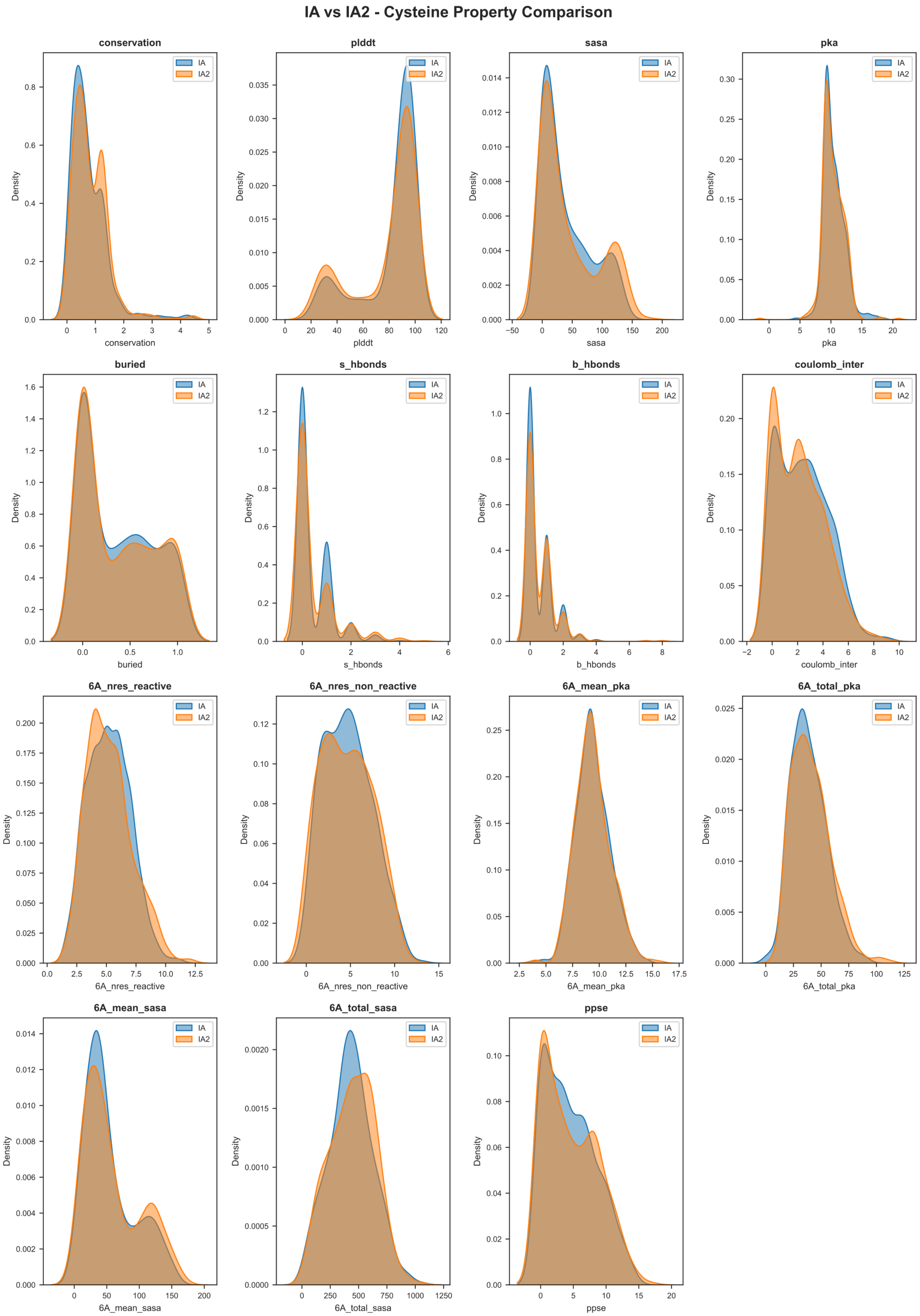
IA vs IA2 - Cysteine Property Comparison
conservation
plddt
sasa
pka
buried
s_hbonds
b_hbonds
coulomb_inter
6A_nres_reactive
6A_nres_non_reactive
6A_mean_pka
6A_total_pka
6A_mean_sasa
6A_total_sasa
ppse
Density
IA
IA2

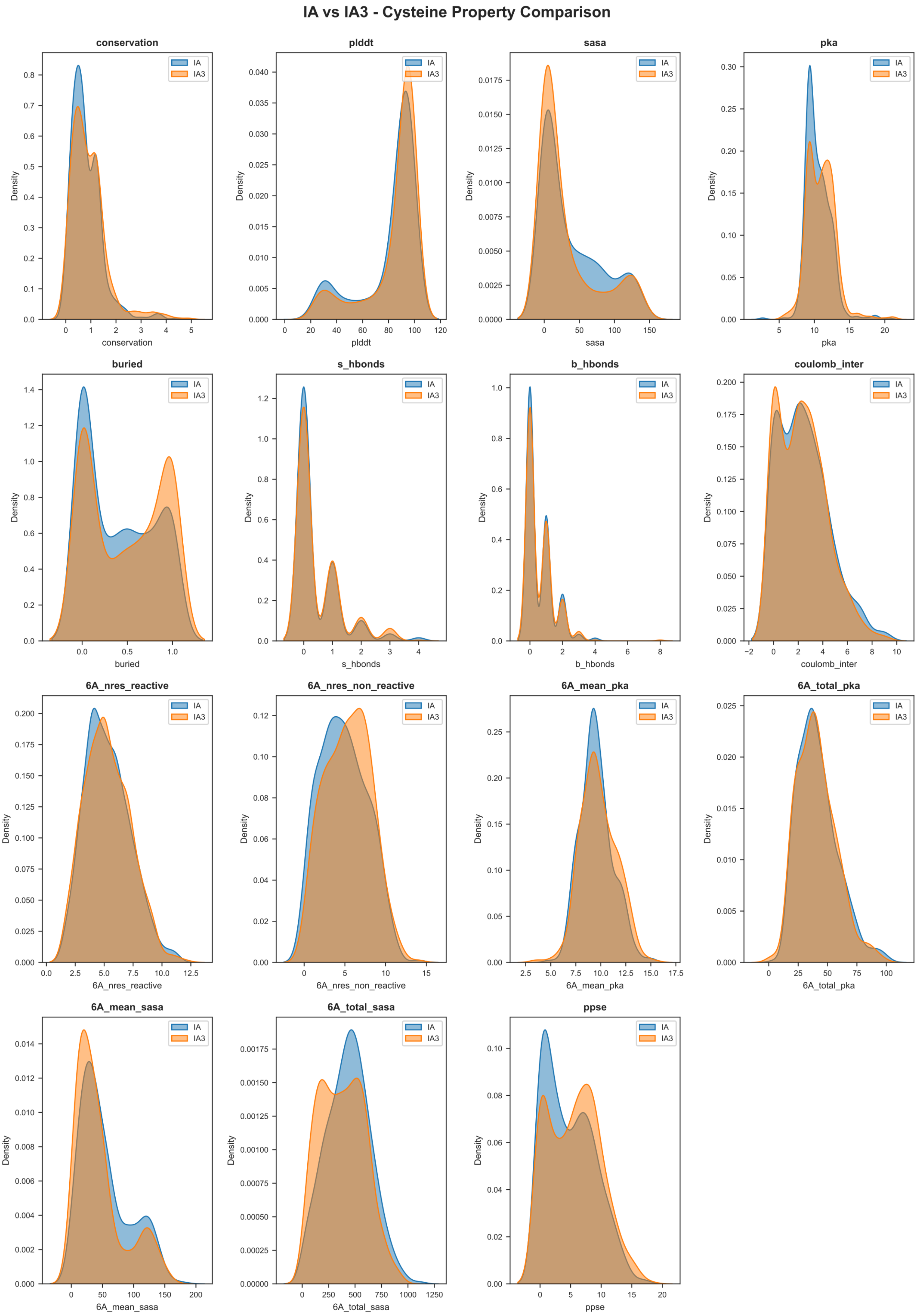
IA vs IA3 - Cysteine Property Comparison
conservation
pldtt
sasa
pka
buried
s_hbonds
b_hbonds
coulomb_inter
6A_nres_reactive
6A_nres_non_reactive
6A_mean_pka
6A_total_pka
6A_mean_sasa
6A_total_sasa
ppse
Density
IA
IA3

### IA vs IA4 - Cysteine Property Comparison

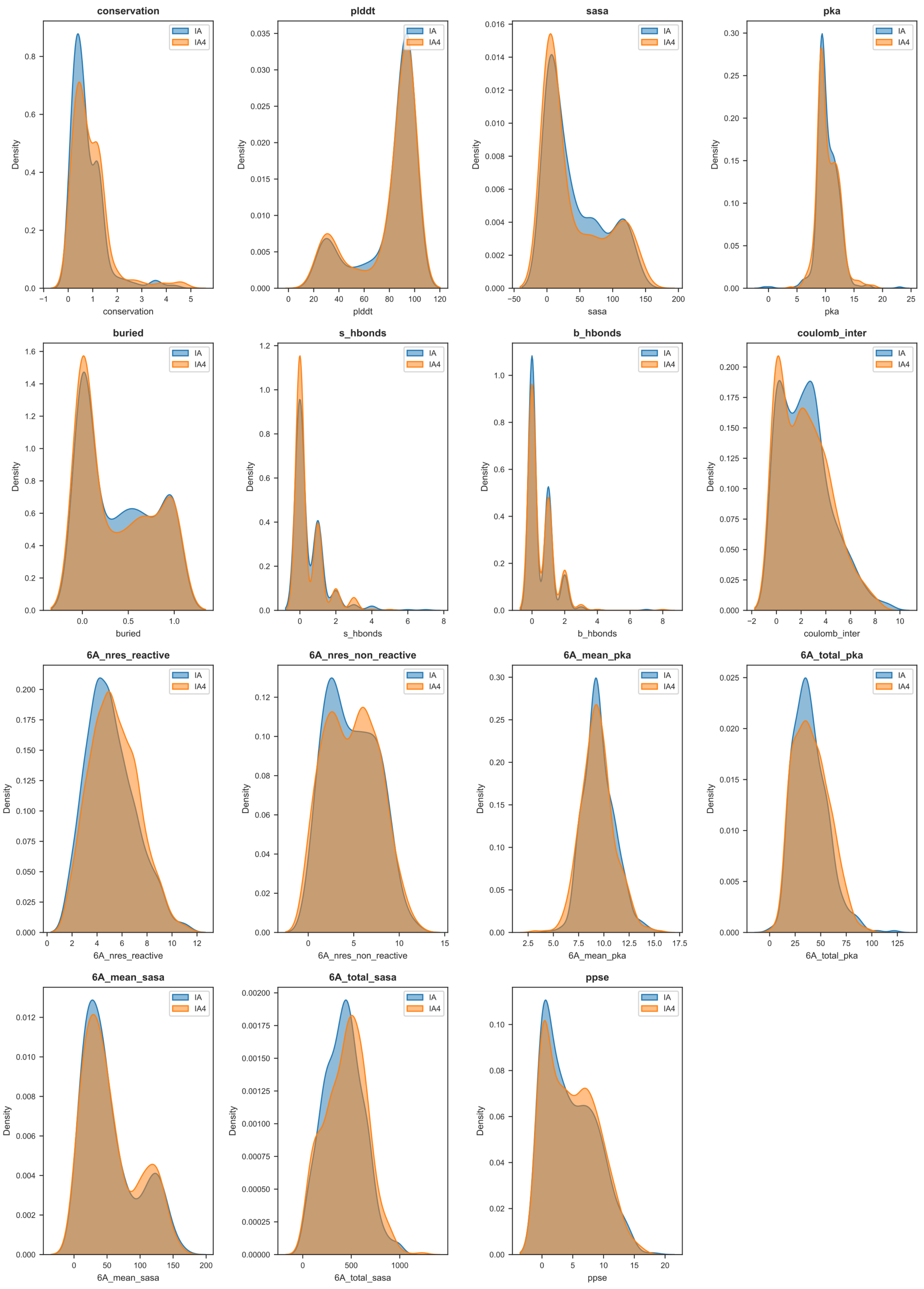

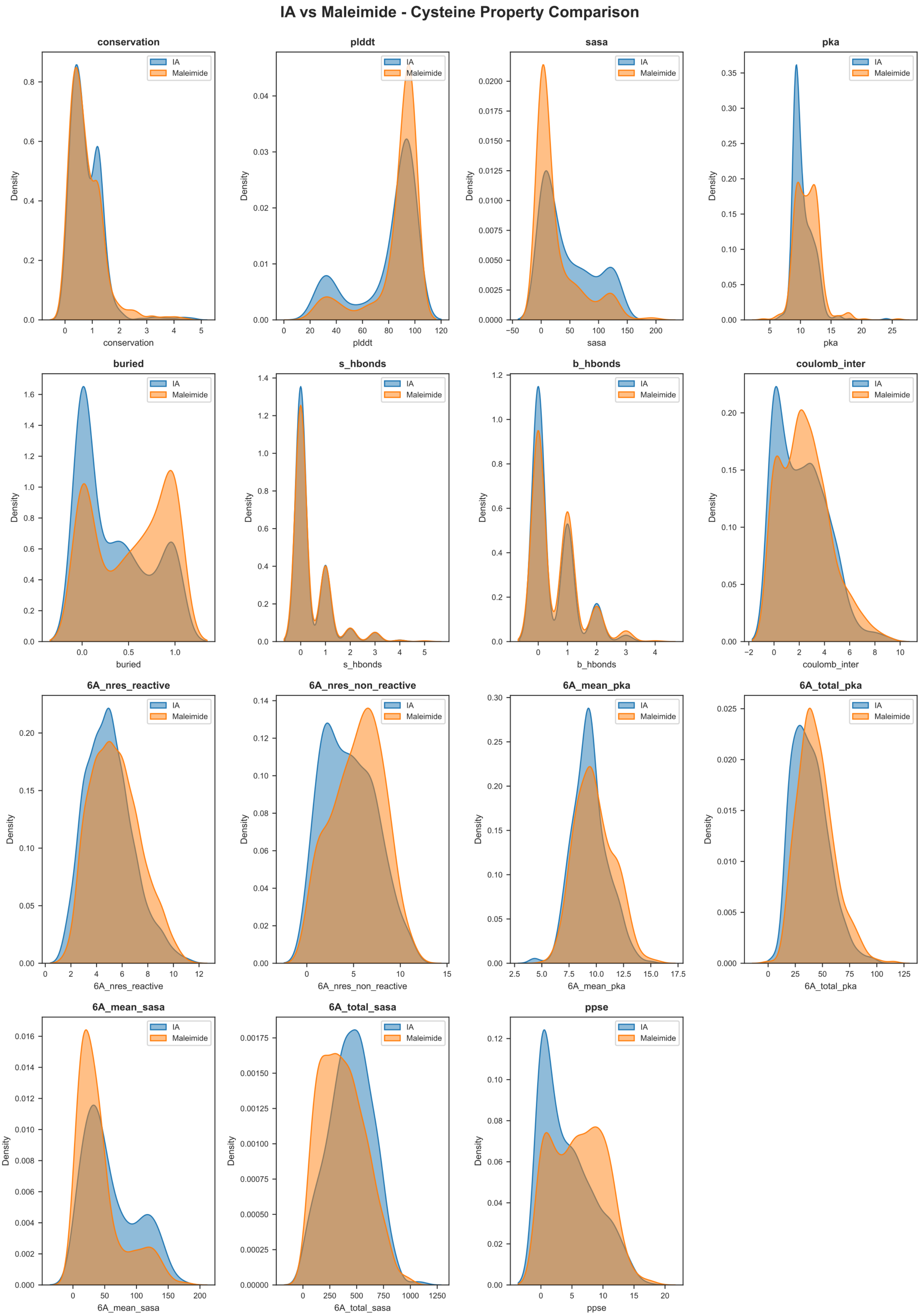
IA vs Maleimide - Cysteine Property Comparison
conservation
plddt
sasa
pka
buried
s_hbonds
b_hbonds
coulomb_inter
6A_nres_reactive
6A_nres_non_reactive
6A_mean_pka
6A_total_pka
6A_mean_sasa
6A_total_sasa
ppse
IA
Maleimide
Density

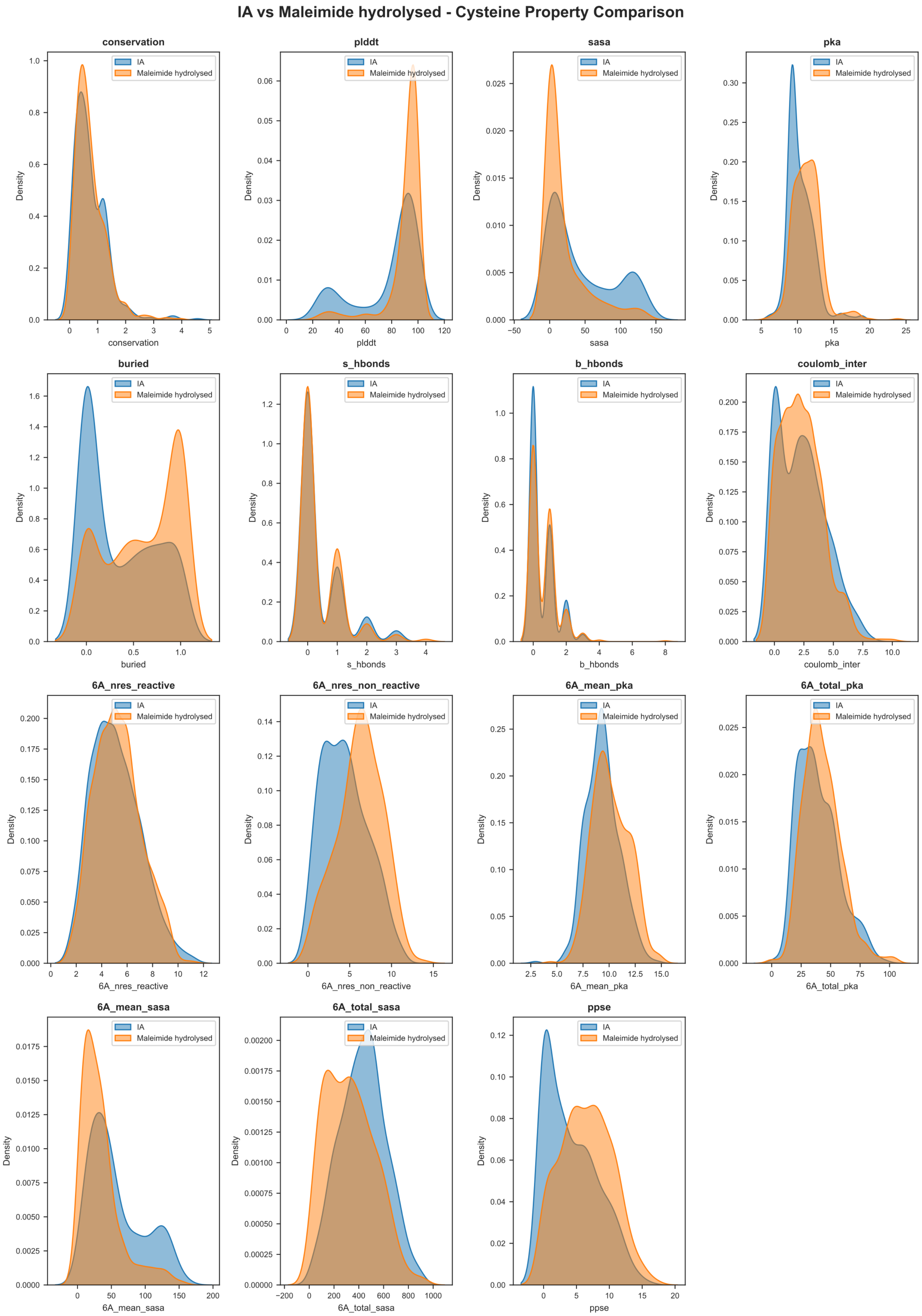
IA vs Maleimide hydrolysed - Cysteine Property Comparison
conservation
plddt
sasa
pka
buried
s_hbonds
b_hbonds
coulomb_inter
6A_nres_reactive
6A_nres_non_reactive
6A_mean_pka
6A_total_pka
6A_mean_sasa
6A_total_sasa
ppse
Density
IA
Maleimide hydrolysed

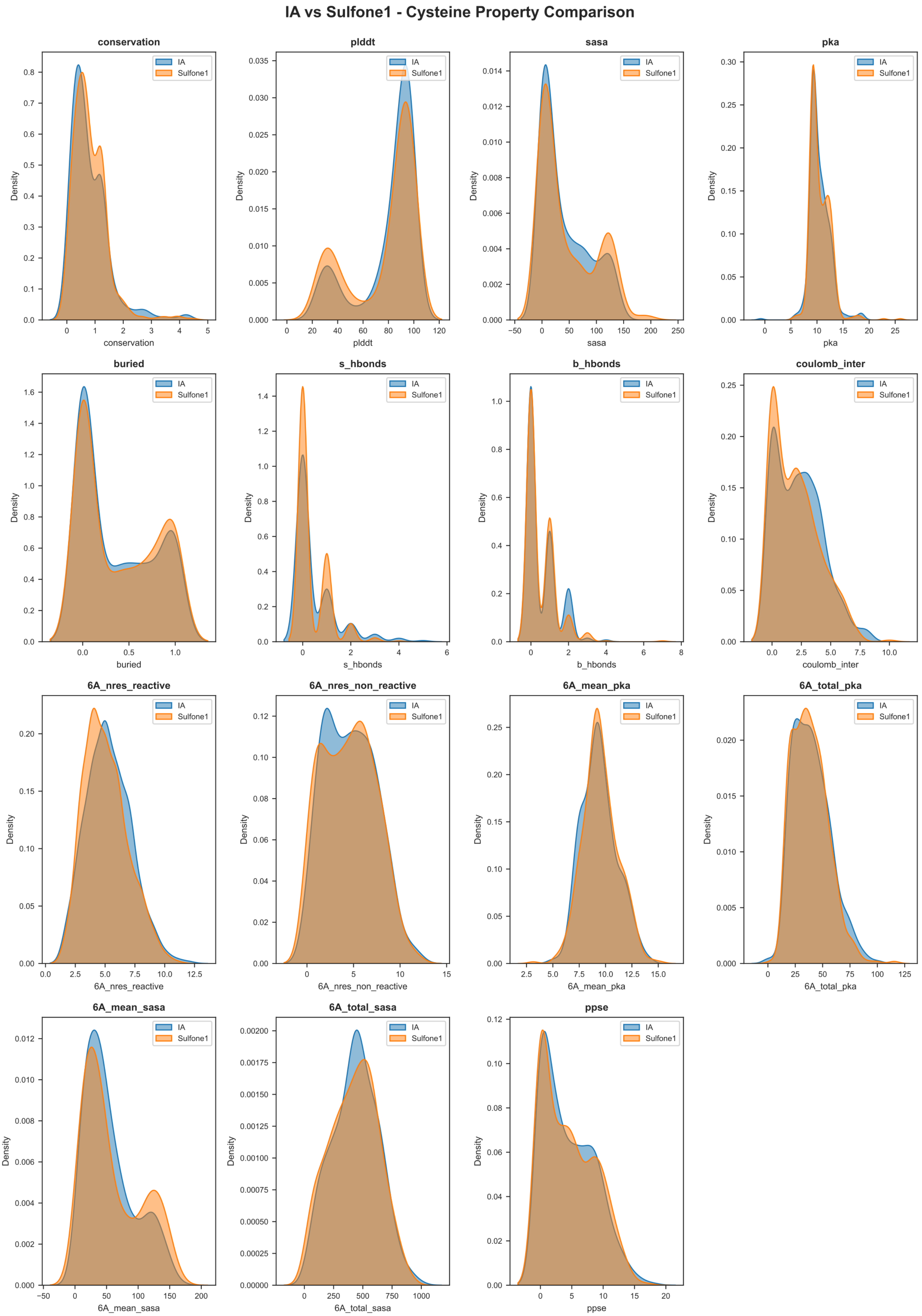
IA vs Sulfone1 - Cysteine Property Comparison
conservation
plddt
sasa
pka
buried
s_hbonds
b_hbonds
coulomb_inter
6A_nres_reactive
6A_nres_non_reactive
6A_mean_pka
6A_total_pka
6A_mean_sasa
6A_total_sasa
ppse
Density
IA
Sulfone1

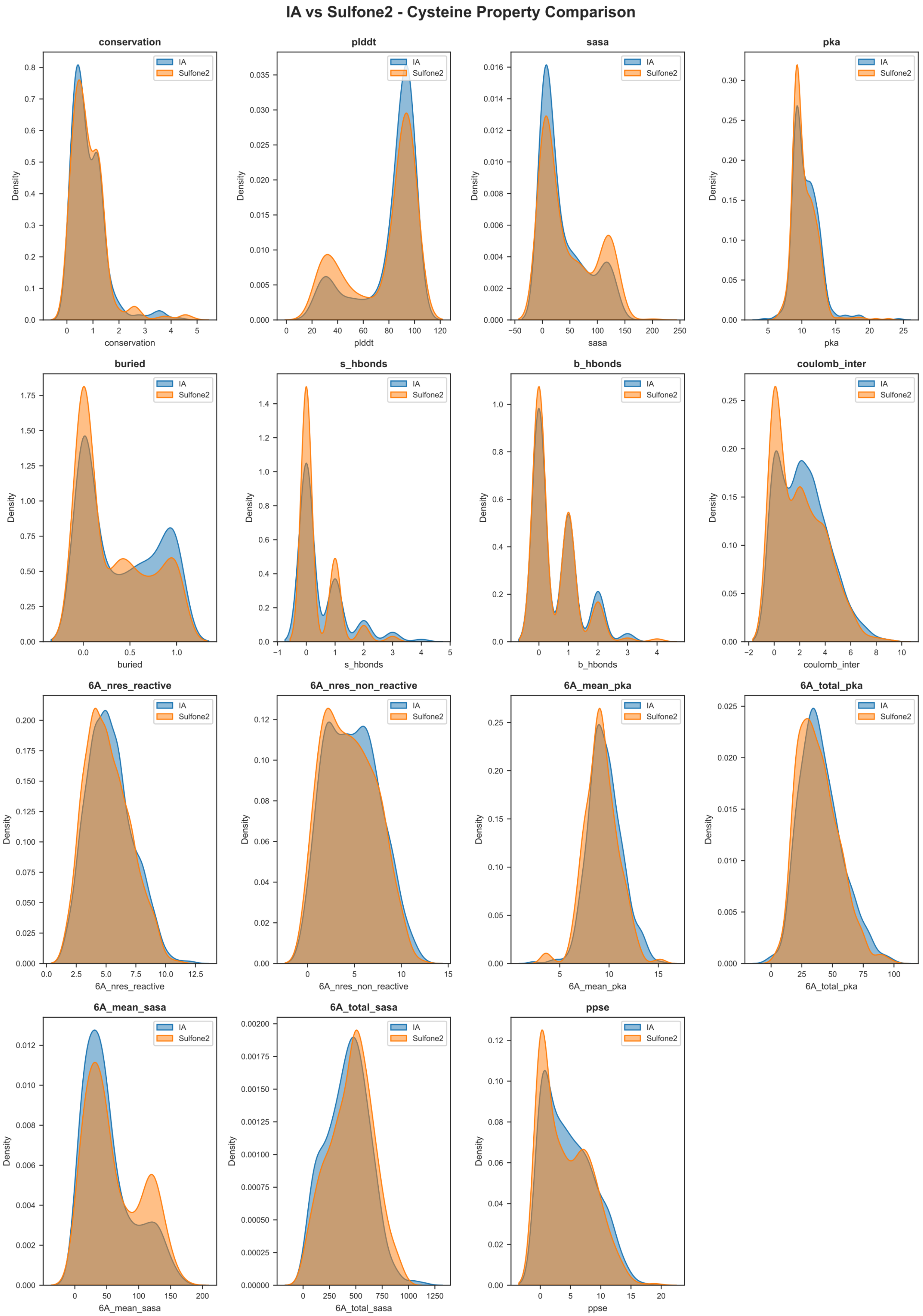
IA vs Sulfone2 - Cysteine Property Comparison
conservation
plddt
sasa
pka
buried
s_hbonds
b_hbonds
coulomb_inter
6A_nres_reactive
6A_nres_non_reactive
6A_mean_pka
6A_total_pka
6A_mean_sasa
6A_total_sasa
ppse
Density
IA
Sulfone2

*Figure S9: Cysteine structural feature distributions in 500 randomly selected, uniquely detected cysteines by IA-DTB vs IA1-DTB, IA2-DTB, IA3-DTB, IA4-DTB, Maleimide2-DTB, Maleimide2 hydrolysed-DTB, Sulfonamide1-DTB and Sulfonamide2-DTB. Distributions are shown as Kernel Density Estimation plots.*

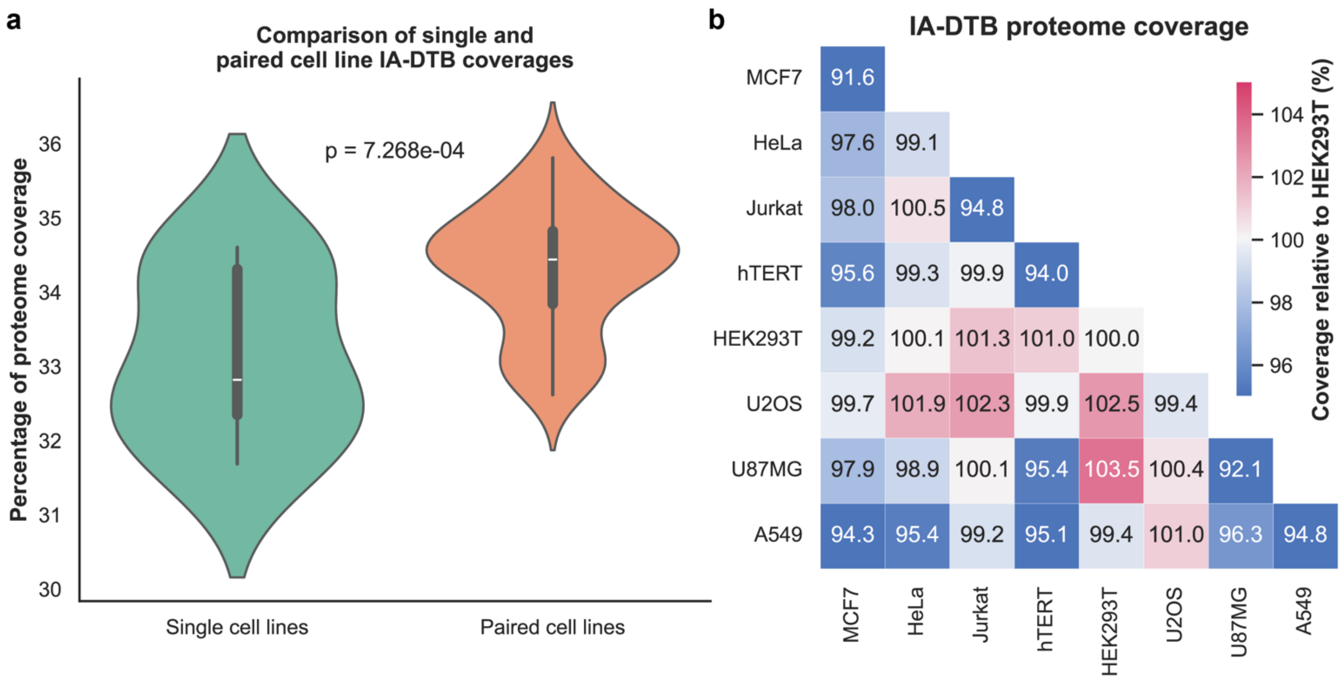


*Figure S10: Mixed lysate IA-DTB proteome coverage. a) Violin plots comparing the percentage of the human proteome covered by IA-DTB in individual cell lines and paired-cell-line mixtures. The displayed p value is from an unpaired two-sided t-test. b Lower-triangular heatmap showing proteome coverage for each cell line and cell-line pair, normalized to the HEK293T single-cell-line coverage, which was set to 100%. Diagonal entries represent individual cell lines; off-diagonal entries represent paired mixtures.*

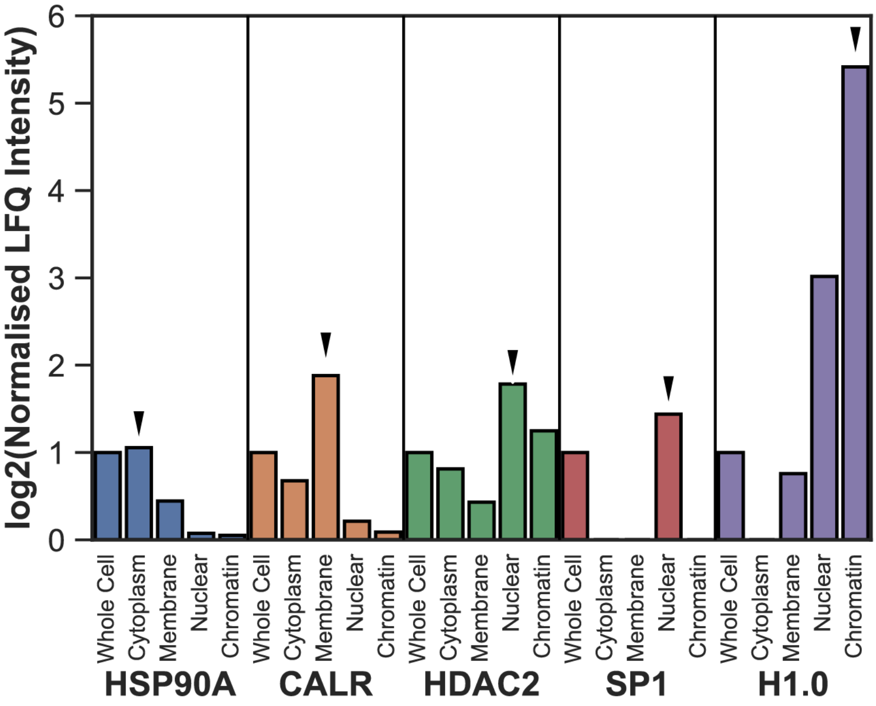


*Figure S11: Quality control of cellular fractionation assay. The expression levels of reference proteins expected to be enriched in specific fractions are shown at each fraction, as indicated by the manufacturer of the assay kit. The expected fraction each protein is expected to be enriched at is indicated with a red arrow.*

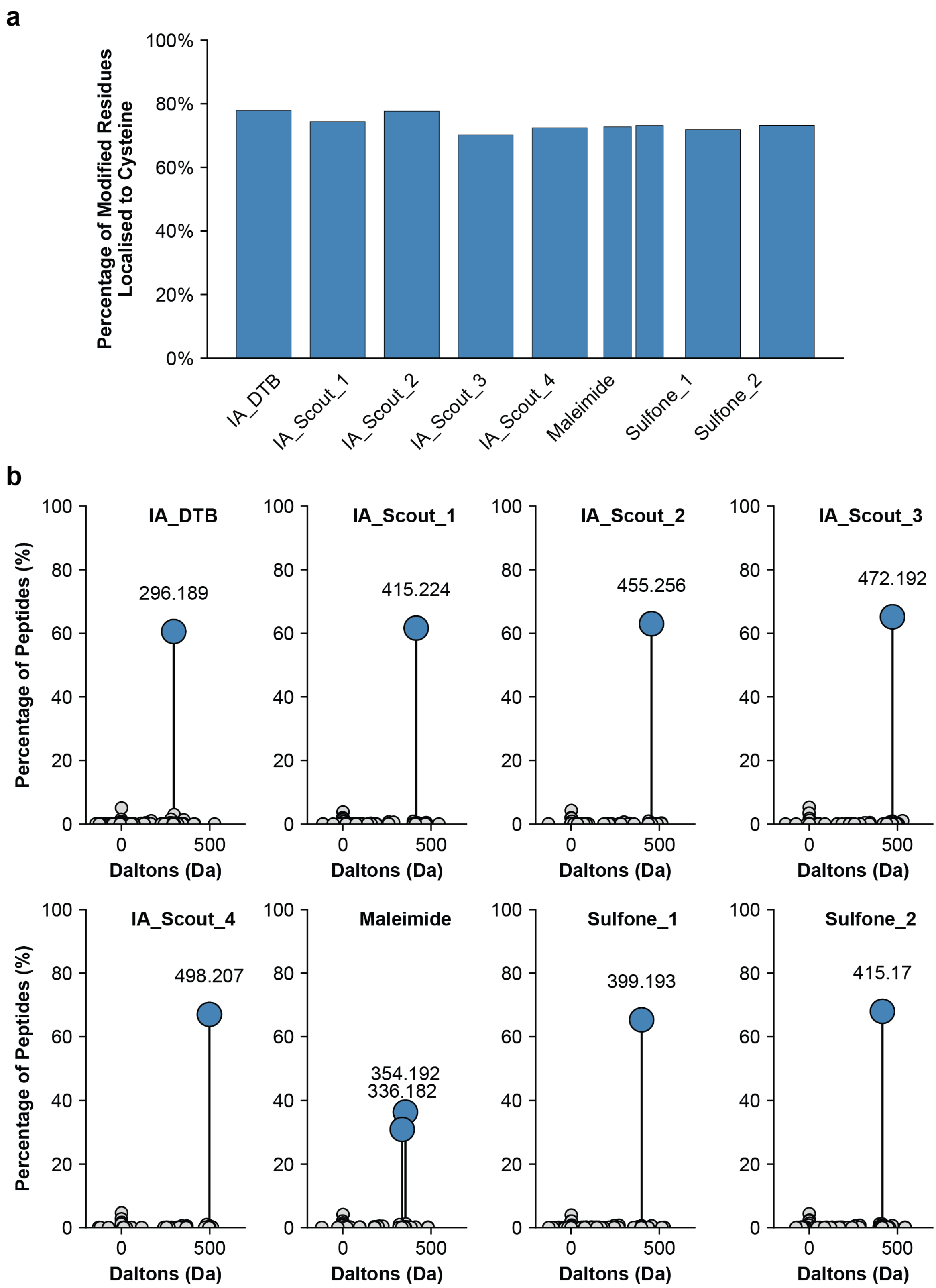


*Figure S12: FragPipe MSFragger open search on the DDA samples of alternative ABPP probes. a) Percentage of modified residues where the ABPP probe PTM was localised to Cysteine, as determined by the FragPipe Open Search. b) All identified masses of modification for each ABPP probe, as determined by the FragPipe Open Search.*
